# Online movements reflect ongoing deliberation

**DOI:** 10.1101/2024.08.19.608669

**Authors:** Jan A. Calalo, Truc T. Ngo, Seth R. Sullivan, Katy Strand, John H. Buggeln, Rakshith Lokesh, Adam M. Roth, Michael J. Carter, Isaac L. Kurtzer, Joshua G.A. Cashaback

## Abstract

From navigating a crowded hallway to skiing down a treacherous hill, humans are constantly making decisions while moving. Insightful past work has provided a glimpse of decision deliberation at the moment of movement onset. Yet it is unknown whether ongoing deliberation can be expressed during movement, following movement onset and prior to any decision. Here we tested the idea that an ongoing deliberation continually influences motor processes—prior to a decision—directing online movements. Over three experiments, we manipulated evidence to influence deliberation during movement. The deliberation process was manipulated by having participants observe evidence in the form of tokens that moved into a left or right target. Supporting our hypothesis we found that lateral hand movements reflected deliberation, prior to a decision. We also found that a deliberation urgency signal, which more heavily weighs later evidence, was fundamental to predicting decisions and explains past movement behaviour in a new light. Our paradigm promotes the expression of ongoing deliberation through movement, providing a powerful new window into understanding the interplay between decision and action.

## INTRODUCTION

When presented with the option of a sweet candy or chocolate, our hand may move back and forth over the two tempting options before we finally make a decision. In this example our online hand movement seems to provide a readout of our ongoing deliberation before a decision. Over the past two decades both behavioural^1,2,3,4^ and neural^5,6^ findings support the idea that deliberation and motor planning are intertwined. Yet it has not been shown that the *ongoing* deliberation—prior to a decision—is expressed throughout online movement execution.

Past work has helped to illuminate the interplay between motor planning and decision-making. During the go-before-you-know paradigm, participants are required to initiate a reaching movement towards multiple potentially correct targets^7,8,1,2,3,4^ At movement onset, participants launched their reaches between or directly at the potentially correct targets. These initial movements reflect priors of the deliberation process, such as representations of the probability of each potential target and movement speed constraints, known during motor planning before movement onset. The correct target is then indicated during the reach via an abrupt and discrete change of evidence (e.g. target colour, phonological input, etc.), where participants would often immediately select and rapidly redirect their movement towards one of the targets. In a different paradigm, humans have similarly been shown to make a “change-of-mind” by rapidly redirecting their movement towards one target^9^ following an initial reach to the other target. These rapid movement redirections were based on evidence provided prior to reaching, demonstrate delayed processing times, and have been interpreted to reflect a second decision. Rapid movement redirections would reflect a final decision, but would obscure a short deliberation and its potential influence on movement. These studies have collectively provided important insights into how priors of deliberation influence motor planning and the timing of midreach decisions, but have not shown that a continuous and ongoing deliberation process directly influences the online movement.

Perceptual decision-making studies manipulate uncertain and continuous evidence, such as the movement of dots^10,11,12,13^ or tokens.^14,15,16^ towards or into potential targets over time, to influence a more prolonged deliberation and subsequent decision. A plethora of work suggests that during deliberation, humans and animals accumulate (integrate) evidence over time to make a decision.^17,18,19,20,11,21,22^ Another competing theory is that an urgency signal increasing over time is multiplied by evidence to cause a decision.^14,15,16,22,23^ A feature of perceptual decision-making tasks is that there is no movement during the deliberation period, a decision is made, and subsequently there is a movement to indicate choice. Thus, even though there is a prolonged deliberation, it does not have the opportunity to be expressed with movement.

Previous studies have collectively provided important insights, but not on how a continuous and ongoing deliberation process directly influences online movement. The goal of this work was to elucidate whether the deliberation process influences online movements, prior to a decision. To investigate we developed a novel paradigm that allows an expression of the ongoing deliberation via movement, prior to a decision. Across three experiments, we permitted movement while concurrently providing uncertain and continuous evidence in the form of 15 tokens that jumped into a left or right target.^14^ In **Experiment 1** we provided participants evidence during posture to test whether the ongoing deliberation can elicit movement onset and subsequently influence online movements, prior to a decision. In **Experiment 2** we provided participants evidence after movement onset, when the motor system was already actively engaged, to determine whether the ongoing deliberation can influence the online movements prior to a decision. In **Experiment 3**, we replicated the results from **Experiment 2** while additionally testing the role of urgency on deliberation. For all experiments we predicted that lateral hand movements would reflect the deliberation process, following movement onset and prior to a decision. Collectively our findings show that the ongoing deliberation, which includes urgency, directly influences online movements.

## RESULTS

### Experimental Design

In **Experiment 1** (n = 17), **Experiment 2** (n = 17), and **Experiment 3** (n = 17), participants made reaching movements while deliberating between two potential targets. For each experiment, there were 15 tokens that moved laterally into either a left or right target (**Fig. 1**). Tokens moved in 160 ms intervals. Each trial was 2400 ms. Participants were instructed to select the target that would finish with the most tokens. They had to make their decision prior to the final token movement. Participants indicated their decision by simultaneously pressing a hand trigger in their non-dominant hand and moving their cursor into their selected target. The hand trigger was crucial in dissociating movements caused by deliberation or a decision. The tokens disappeared once participants pressed the hand trigger to prevent the participants from changing their decision with later evidence. Critically, participants were free to move their hand laterally during each trial, allowing us to measure whether deliberation—prior to a final decision—influenced movement.

**Figure. 1:**
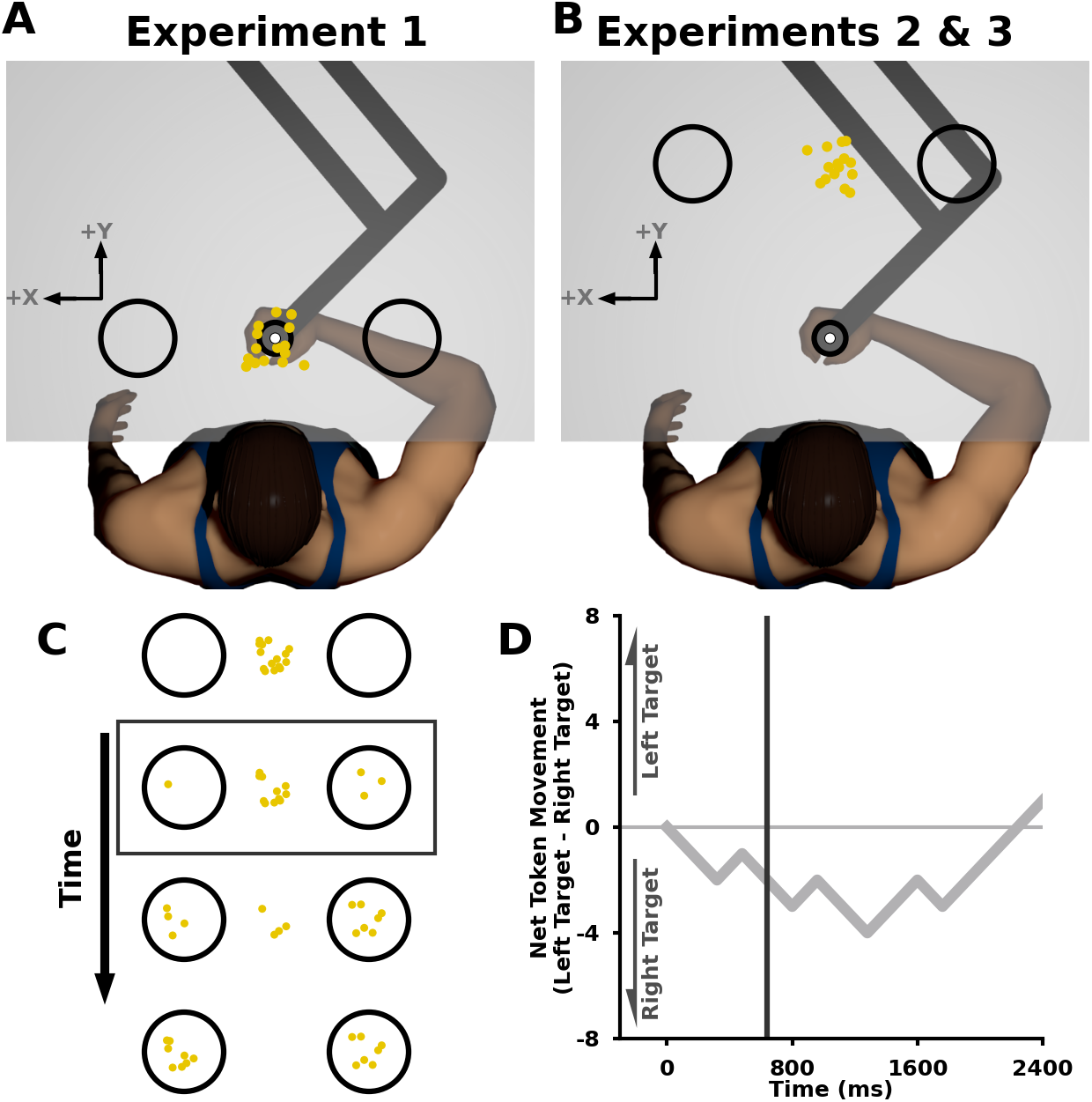
Experimental Design. **A**,**B)** Participants grasped a robotic manipulandum (Kinarm) using their dominant hand and a hand trigger in their non-dominant hand. A semi-silvered mirror projected images from an LCD screen above. A cursor (white circle) represented their hand position. **A)** In **Experiment 1**, participants began with the cursor within a start position (small black circle) between two targets (large black circles) that were 20 cm to the left and right of the start position. After 400ms in the start position, the trial would begin and fifteen tokens would appear (yellow circles) between the two targets. The tokens moved individually into the left or right target over time. Participants were instructed to select the target which would finish with the most tokens as soon as they were confident. They indicated their decision by simultaneously pressing the hand trigger in their non-dominant hand and moving the cursor into the corresponding target. **B)** In **Experiments 2** and **Experiment 3**, the targets were placed 30 cm forward of the start position as well as 20 cm to the left and right. Tokens began moving once participants left the start position. Participants were also instructed not to stop or move backwards. **C)** An example of the participant display while the tokens moved into the left or right target over time (y-axis). **D)** Net token movement (left target - right target tokens, y-axis) over time (x-axis) of an example token pattern. The dark grey box in (**C**) and the dark grey vertical line (**D**) correspond to the same time point.

The goal of **Experiment 1** was to determine if ongoing deliberation can elicit and subsequently influence movements, prior to a final decision, when evidence was initiated during posture. The targets were placed on the right and left side of the start position (**Fig. 1A**). The trial began after participants held their hand within a 2 cm wide start position for 400 ms. Participants experienced 216 randomly interleaved trials consisting of pseudo-random token patterns and bias token patterns (See **Methods, Supplementary A**). The bias token patterns allowed us to probe how controlled patterns of evidence influenced deliberation and consequently movement. During the bias token patterns the first three tokens moved individually into the left or right target (i.e., left bias or right bias), the next three tokens moved individually into the opposite target, and the remaining tokens moved with an 80% probability into the left or right target (i.e., left target or right target; **Fig. 2A-D**).

**Figure. 2:**
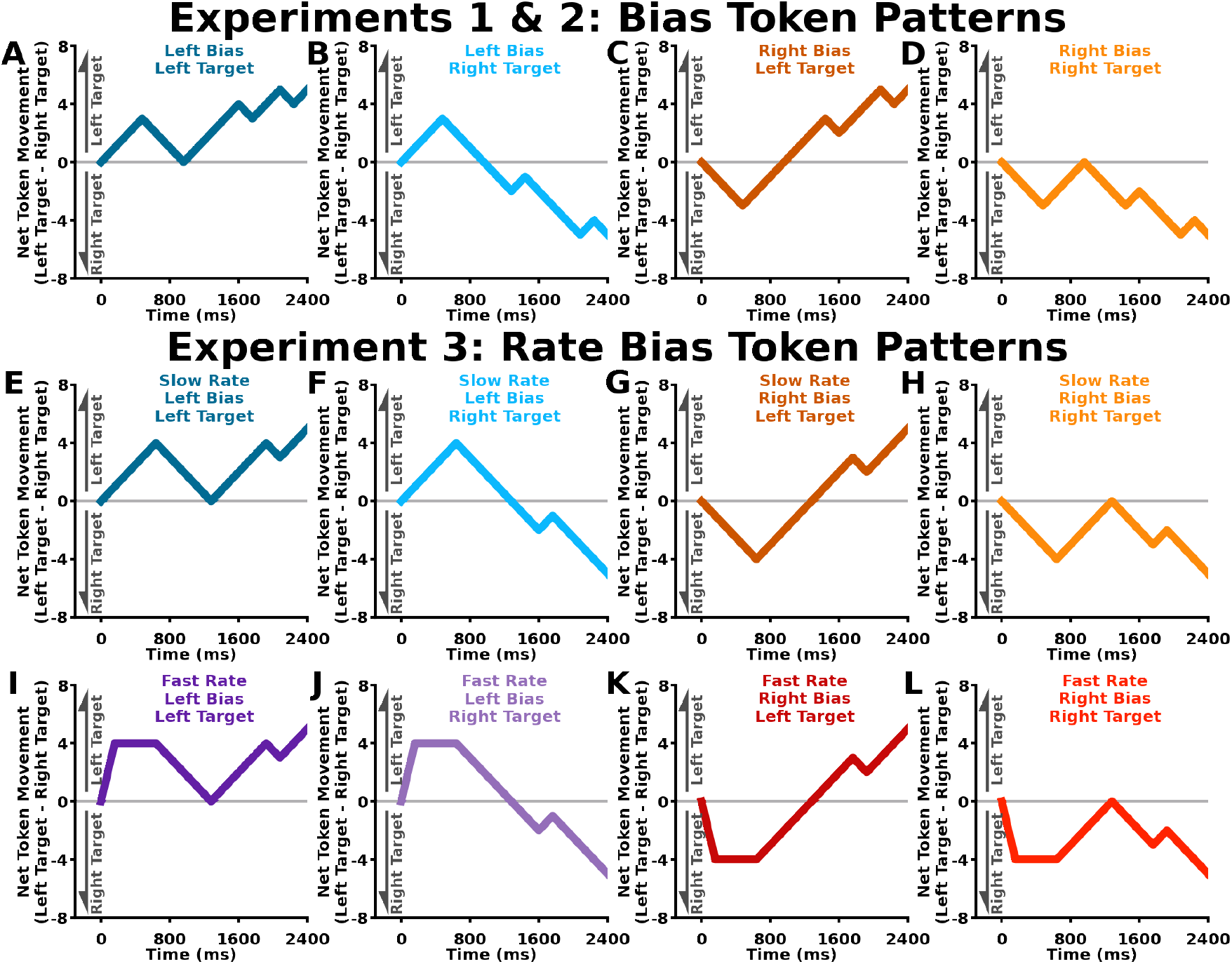
Bias Token Patterns. **A-L)** Net Token Movement (left target - right target tokens, y-axis) over time (x-axis) for sample bias token patterns. **A-D)** Bias token patterns in **Experiment 1** and **Experiment 2. A-D)** Bias token patterns: the first three tokens moved individually into the left or right “bias” target, the next three tokens moved individually into the opposite target, and then the remaining tokens moved with an 80% probability into the left or right target. **E-L)** Rate bias token patterns in **Experiment 3. E-H)** Slow rate bias token patterns: the first four tokens moved individually into the left or right “bias” target, the next four tokens moved individually into the opposite target, and then the remaining tokens moved with an 80% probability into the left or right target. **I-L)** For the fast rate bias token patterns: the first four tokens moved together into the left or right “bias” target at 160 ms after the beginning of the trial. No other tokens moved until 800 ms after the beginning of the trial. The slow rate and fast bias token patterns were identical past 800 ms after the beginning of the trial. For each experiment, the bias token patterns were interleaved with psuedorandom token patterns.

The goal of **Experiment 2** was to determine if ongoing deliberation was reflected in movements, prior to a final decision, after movement onset when the motor system was already actively engaged. In this experiment, the targets were placed forward and either side relative to the start position (**Fig. 1B**). To actively engage the motor system, the trial began when participants moved forward out of the start position. Similar to others,^3,4^ participants were instructed to not stop moving forward after leaving the start position. **Experiment 2** used the same token patterns as **Experiment 1**.

The goal of **Experiment 3** was to replicate the results found in **Experiment 2** while also elucidating the roles of evidence accumulation or urgency on deliberation and consequent movement. **Experiment 3** was the same as **Experiment 2**, except we used different bias token patterns. In **Experiment 3**, participants experienced 336 randomly interleaved trials consisting of pseudorandom token patterns, slow rate bias token patterns and fast rate bias token patterns.

In the slow rate bias token patterns, the first four tokens moved individually into the left or right bias target, the next four tokens moved individually into the opposite target and the remaining tokens moved with an 80% probability into the left or right target (**Fig 2E-H**). The fast rate bias token patterns were identical to the slow rate bias token pattern except the first 4 tokens moved at once into the corresponding bias target (**Fig. 2I-K**). Critically, the slow rate and fast rate token patterns lead to unique decision times depending on how humans accumulate evidence and or rely on urgency during deliberation.

### Individual Movement Behaviour

We were primarily interested in the lateral hand position at the estimated decision time. Lateral hand position at estimated decision time provided a measure of the influence of ongoing deliberation on the movement. In other words, the lateral hand position at estimated decision time precludes movement that is a result of a final decision and subsequent action. Estimated decision time was calculated by subtracting a neural plus mechanical delay from the trigger time on each trial (**see Methods**;**Fig. 3A,B**). We examined lateral hand position at estimated decision time to compare between conditions (**Fig. 3C**).

**Figure. 3:**
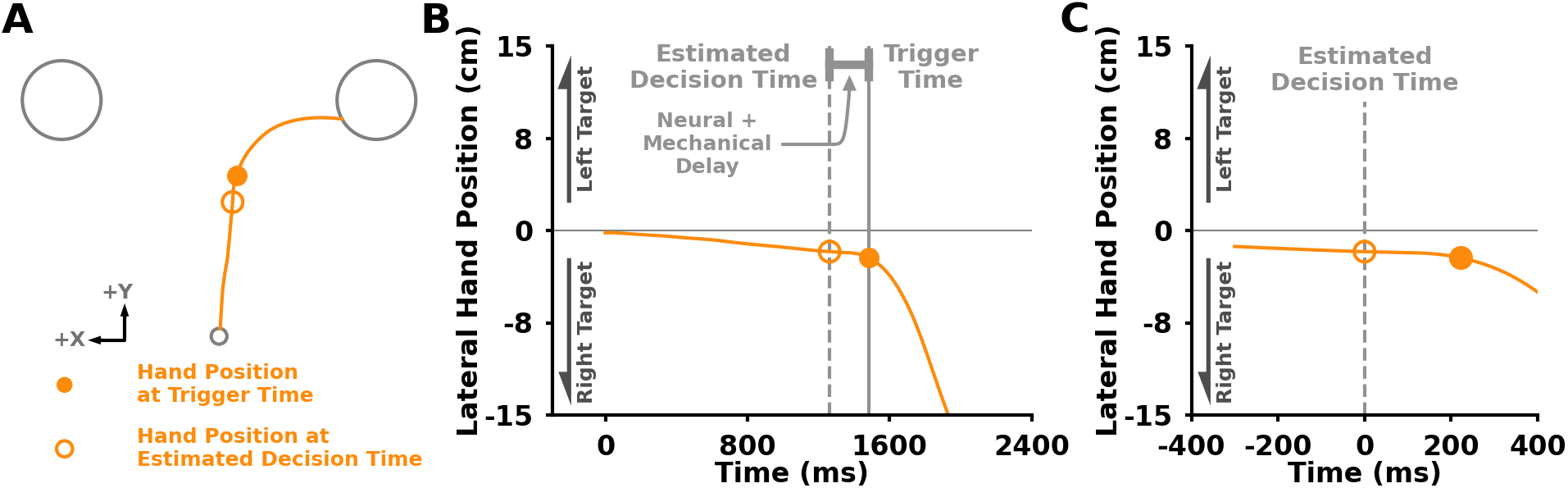
Analysis Description. **A)** Hand path for a single example trial in **Experiment 2**. Solid circles represent the hand position when the participant pressed the hand trigger (trigger time). Empty circle represents the hand position at estimated decision time. Estimated decision time was calculated by subtracting a neural and mechanical delay from the trigger time on a trial-by-trial basis. Neural + mechanical delay was estimated for each participant using a reaction time task (see **Supplementary A**). **B)** Lateral hand movement (y-axis) over time (x-axis). Solid grey line represents when the hand trigger was pressed. Dashed grey lane represents estimated decision time. **C)** Lateral hand position (y-axis) over time (x-axis) aligned to estimated decision time. The lateral hand position at the estimated decision time allows us to look at the influence of deliberation on movement, prior to a final decision.

**Figure 4** presents results by representative individuals in each experiment. In **Experiment 1**, this participant did not initiate lateral movements prior to their estimated decision time (**Fig 4A-D**). In **Experiment 2**, the participant displayed lateral movements aligned with token bias direction prior to the estimated decision time (**Fig. 4E-H**), which reflects movement that occurred before their final decision. Moreover, their lateral hand position aligned with the token bias direction (**Fig. 4H**). In **Experiment 3**, the representative participant displayed lateral movements that aligned with the direction of the bias in both slow rate bias (**Fig. 4I-L**) and fast rate bias (**Fig. 4M-P**) token patterns. That is, the displayed participants in **Experiments 2** and **3** moved with the evidence prior to a final decision, suggesting that their movements were influenced by the ongoing deliberation.

### Group Movement Behaviour

**Figure 5** displays the average group movement behaviour for the three experiments. We predicted that the lateral hand movements would be influenced by the ongoing deliberation, prior to a decision. For example, a participant that is considering the left target will move towards the left target, prior to their final decision. We show the average lateral hand trajectories over time for **Experiment 1, 2**, and **3** (**Fig. 5A,D,G,J**). However, it is important to examine lateral hand positions at the estimated time (**Fig. 5B,E,H,K**), which reflects movement caused by deliberation prior to a final decision.

#### Hand movements are influenced by deliberation when the motor system is actively engaged

In **Experiment 1**, lateral hand position at the estimated decision time was not impacted by the token patterns (**Fig. 5B**). We did not find a significant main effect of bias [F(1,16) = 3.681, p = 0.073], main effect of target [F(1,16) =1.016, p = 0.328], or an interaction between bias and target [F(1,16) = 0.067, p = 0.799] on lateral hand position at estimated decision time (**Fig. 5C**). The results in **Experiment 1** do not support the idea that the deliberation process continuously interacts with the motor control processes to influence online movements, specifically when evidence is initially presented while in posture. In **Experiment 2**, we examined the influence of ongoing deliberation on the motor control system when the motor system was actively engaged. Here participants displayed lateral hand positions at estimated decision time that was aligned with the direction of the token bias (**Fig. 5E**). Specifically, we found a significant main effect of bias [F(1,16) = 11.533 p = 0.004] on lateral hand position at estimated decision time. We did not find an interaction between bias and target [F(1,16) = 0.300, p = 0.591] nor a main effect of target [F(1,16) = 0.255, p = 0.620]. When collapsing across target, as expected we found significantly different lateral hand positions at estimated decision time between left bias and right bias token patterns (**Fig. 5F**;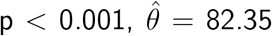). Moveover, our findings and interpretation were consistent when we very conservatively looked further back in time (See **Supplementary B**), along with pseudorandom token patterns (e.g., 20%, 35%, 50%, 75%, and 80% left target probability; see **Supplementary C**). The findings in **Experiment 2** support the hypothesis that the ongoing deliberation process influences online movements, prior to a decision, when the motor system is actively engaged.

**Figure. 4:**
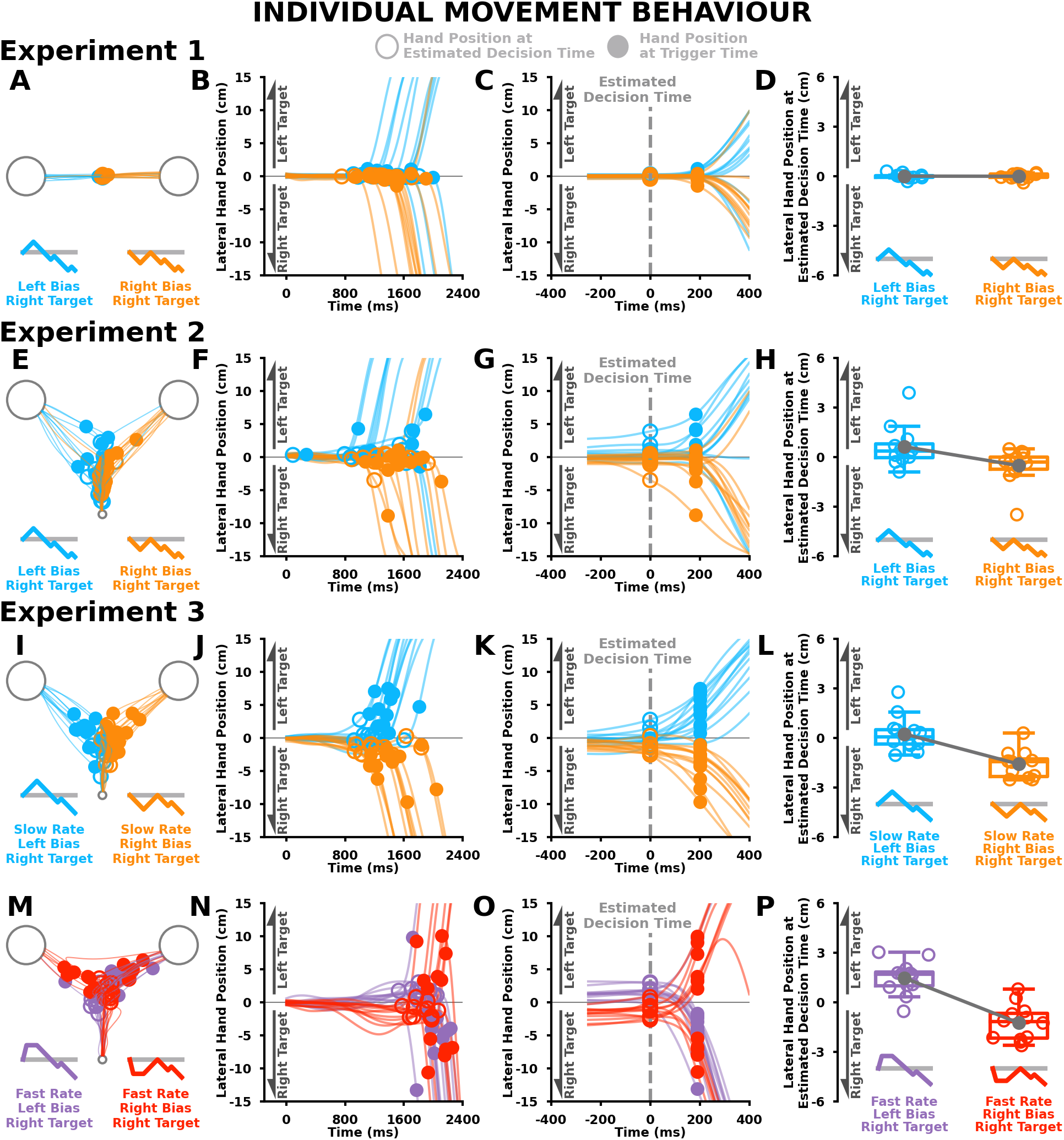
Individual Movement Behaviour. **A-H)** Individual participant movement behaviour for left bias, right target (light blue) and right bias, right target (light orange) token patterns in **Experiment 1** (**A-D**) and **Experiment 2** (**E-H**). **I-P)** Individual participant movement behaviour in **Experiment 3** for **I-L)** slow rate, left bias, right target (light blue) and slow rate, right bias, right target (light orange) token patterns. **M-P)** Fast rate, left bias, right target (light purple) and fast rate, right bias, right target (light red) token patterns. Solid circles represent hand position at trigger time. Empty circles represent hand position at estimated decision time. **A,E,I,M)** Individual participant reaching trajectories. **B,F,J,N)** Individual participant lateral hand positions (y-axis) over time (x-axis). **C,G,K,O)** Individual participant lateral hand positions (y-axis) over time (x-axis) time aligned to estimated decision time. Vertical grey dashed line at 0 ms represents estimated decision time. **D,H,L,P)** Individual participant lateral hand positions at estimated decision time (y-axis) between bias token patterns (x-axis). In **Experiment 1**, this participant did not display differences in lateral hand position at estimated decision time between conditions. Participants in **Experiment 2** and **Experiment 3** show differences in lateral hand positions at estimated decision time between left and right bias conditions.

**Figure. 5:**
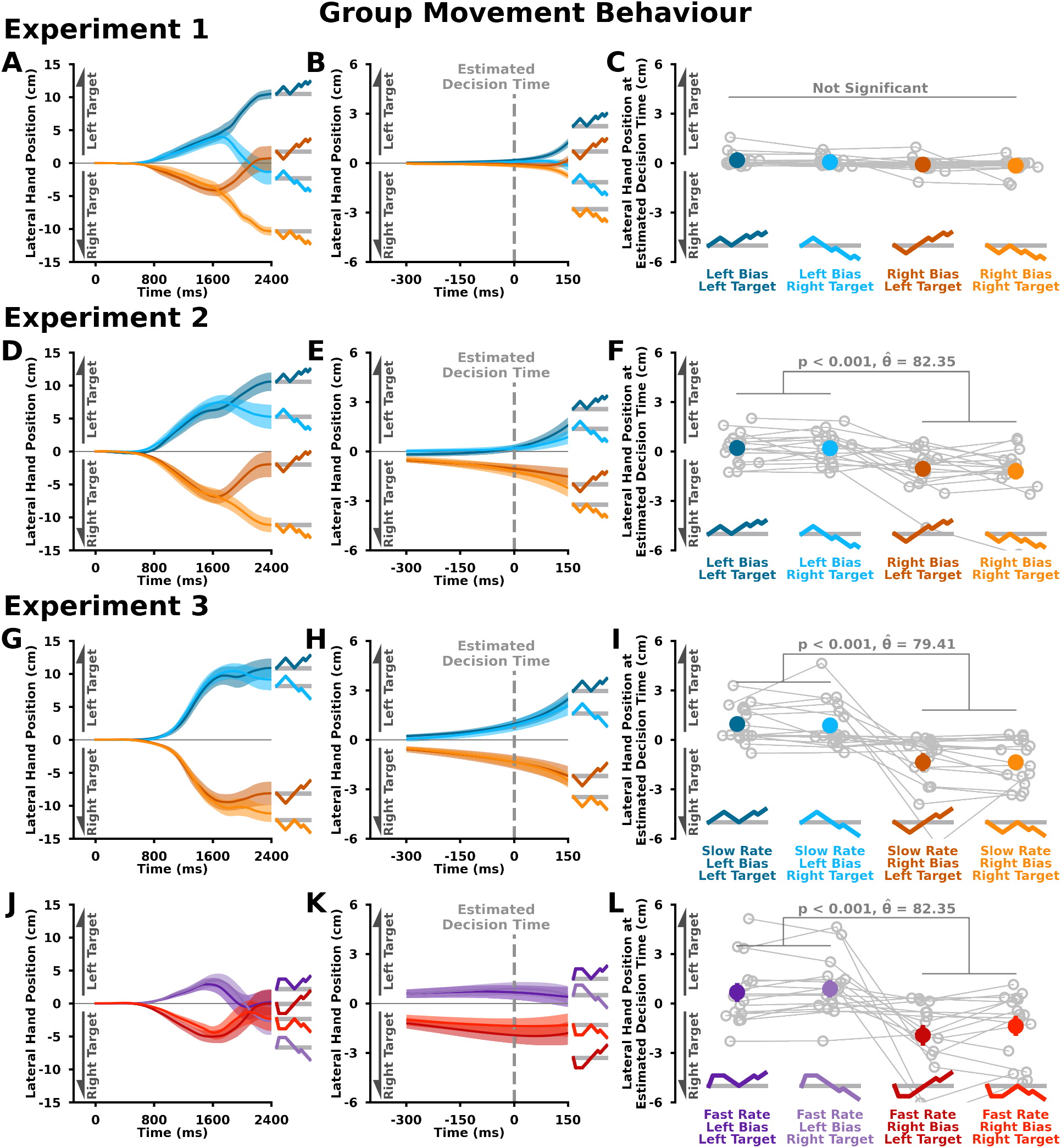
Group Movement Behaviour. **A-F)** Average participant movement behaviour for bias token patterns in **Experiment 1** (**A-C**) and **Experiment 2** (**D-F**). **G-L)** Average participant movement behaviour in **Experiment 3** for **G-I)** slow rate bias token patterns and **J-L)** fast rate bias token patterns. Solid lines represent group mean trajectories for each condition. Shaded regions represent ±1 standard error. **A,D,G,J)** Average participant lateral hand positions (y-axis) over time (x-axis). **B,E,H,K)** Average participant lateral hand positions (y-ax is) over time (x-axis) time aligned to estimated decision time. Vertical grey dashed line at 0 ms represents estimated decision time. **C,F,I,L)** Average participant lateral hand positions at estimated decision time (y-axis) across bias token patterns (x-axis). In **Experiment 1**, there were no significant differences in lateral hand positions at estimated decision time between bias token patterns. Participants in **Experiment 2** and **Experiment 3** were significantly more towards the left target in left bias token patterns compared to right bias token patterns at the estimated decision time (p<0.001 for all comparisons). Taken together, these results suggest that lateral hand movements reflect the ongoing deliberation during movement prior to a decision.

In **Experiment 3** we replicated the movement behaviour findings of **Experiment 2**. We analyzed lateral hand position at estimated decision times separately for slow rate and fast rate token patterns, since they had different decision times (see **Group Decision-Making Behaviour** below). For the slow rate token patterns we found a significant main effect of bias [F(1,16) = 14.663, p = 0.001] on lateral hand position at estimated decision time, but no main effect of target [F(1,16) = 0.0875, p = 0.771] or bias and target interaction [F(1,16) = 0.040, p = 0.844]. For the fast rate token patterns we found a significant main effect of bias [F(1,16) = 9.114, p = 0.008] and a significant main effect of target [F(1,16) = 4.834, p = 0.043] on lateral hand position at estimated decision time, and not a bias and target interaction [F(1,16) = 1.297, p = 0.272]. We found significantly different lateral hand position at estimated decision time between left bias and right bias conditions for both slow rate bias token patterns (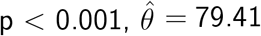, **Fig. 5I**) and the fast rate bias token patterns (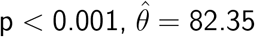, **Fig. 5L**). Again, differences in lateral hand position support the hypothesis that ongoing deliberation influences movement, prior to a decision, when the motor system is actively engaged.

Taken together, our results from **Experiments 1, 2**, and **3** support the idea that the ongoing deliberation process influences hand movement—prior to a decision—when the motor system is actively engaged but not during posture.

### Group Decision-Making Behaviour

#### Humans relied less on early evidence when making decisions

We were also interested in the processes that underscore the deliberation. **Figure 6** shows the estimated decision times for each bias token pattern and experiment. In **Experiment 1**, we found a significant main effect of bias [F(1,16) = 7.222, p = 0.016] on estimated decision time, but there were no significant differences in followup mean comparisons (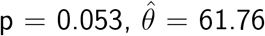, **Fig. 6A**). We did not find a significant main effect of target [F(1,16) = 0.606, p = 0.447] or an interaction between bias and target [F(1,16) = 0.930, p = 0.349] on estimated decision time. In **Experiment 2**, we did not find a significant main effect of bias [F(1,16) =0.989, p = 0.335], significant main effect of target [F(1,16) < 0.001, p = 0.993], or an interaction between bias and target [F(1,16) = 0.154, p = 0.700] on estimated decision time (**Fig. 6B**). Interestingly, participants made faster decisions during **Experiment 2** compared to **Experiment 1** (p < 0.003, *hatθ* = 67.76). One possibility for our result is that decisions are made faster when the motor system is actively engaged, supporting bidirectional interactions between decision and motor processes. In **Experiment 3**, we found a significant main effect of rate [F(1,16) = 27.18, p < 0.01] on estimated decision time (**Fig. 6C**). Counterintuitively, we found that participants made earlier decisions in slow rate compared to fast rate token patterns (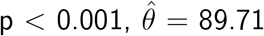, **Fig. 6C,7A**). We did not find main effects of target [F(1,16) = 0.689, p = 0.419], main effect of bias [F(1,16) = 0.588, p = 0.454], nor any significant interactions (p > 0.105). The selection rates for each token pattern are shown in **Supplementary D**.

**Figure. 6:**
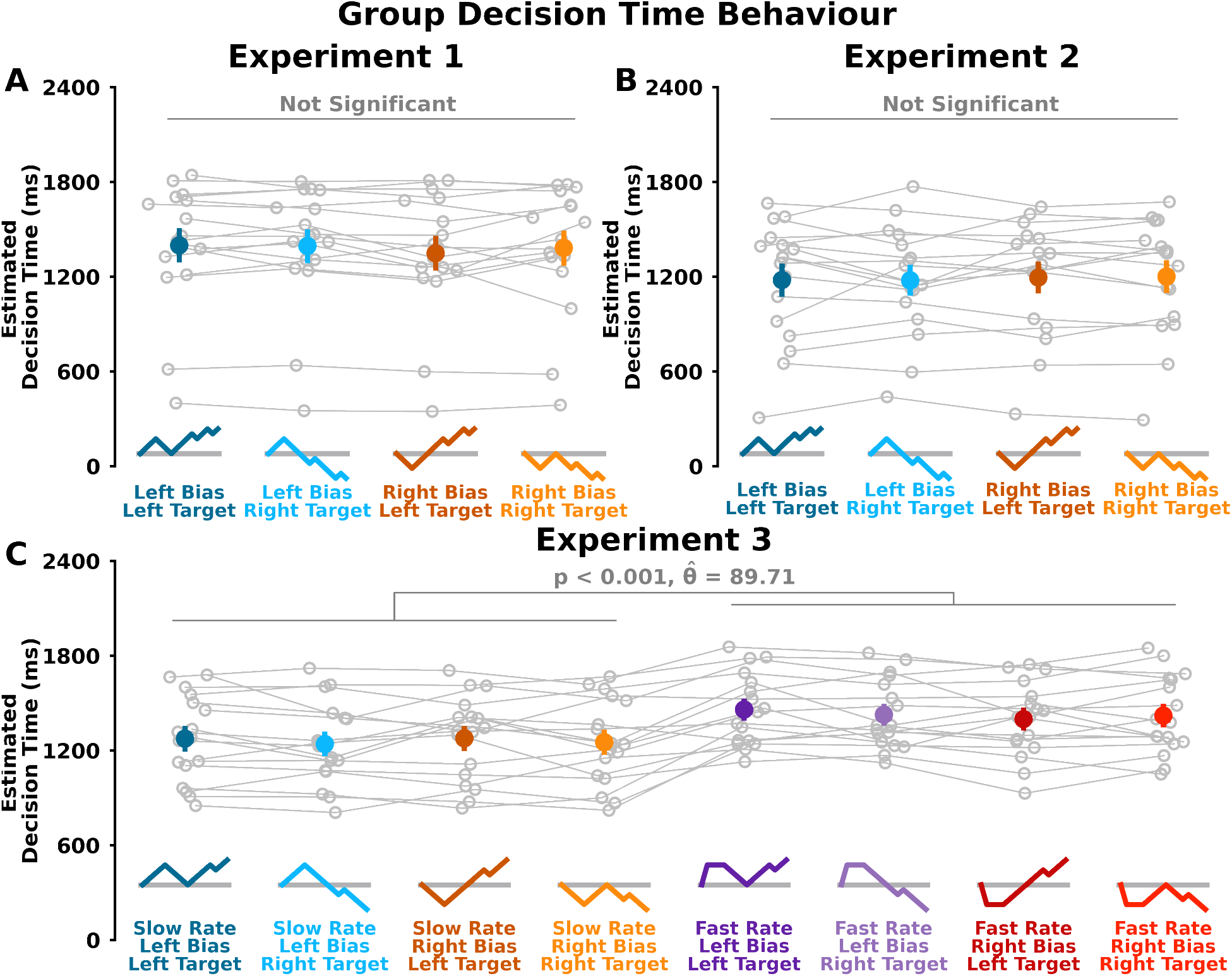
Group Decision Time Behaviour. Estimated decision time (ms; y-axis) in **A) Experiment 1, B) Experiment 2**, and **C) Experiment 3** across bias token patterns (x-axis). Open grey circles and connected grey lines represent individual participants. Closed coloured circles (and error bars) represent mean (and standard error of the mean) for each token pattern. Estimated decision time did not change between bias token patterns in **A) Experiment 1** (p > 0.05) and **B) Experiment 2** (p > 0.05). **C)** Participants had earlier estimated decision times in **Experiment 3** for slow rate token patterns (blue and orange colours) compared to fast rate token patterns (purple and red colours; p < 0.001), suggesting a greater temporal weighing of later evidence in the decision-making process.

Above we did not find a significant bias and target interaction on estimated decision time. This pattern is consistent with past work by Cisek (2009) that proposed that urgency is involved with deliberation. As a reminder, urgency represents less reliance on early evidence compared to later evidence when making a decision. Interestingly and counterintuitively, we found that participants made earlier decisions with a slow rate token pattern compared to the fast rate token pattern. This finding strongly align with the idea that decision making processes more heavily value information that is presented later in time (i.e., second, third and fourth tokens in the slow rate token pattern) compared to the same information presented earlier in time (i.e., second, third and fourth tokens presented earlier in time during the fast rate token pattern). However, as shown below in **Decision-making models**, the presence of both urgency and evidence integration best explain the reported estimated decision times.

### Computational Modelling

Our central focus was to investigate the interaction between the decision-making and motor control processes. To this end, we used a computational framework that combines a decision-making model and an optimal feedback control model.

#### Decision-making models

Before combining decision and motor models, we first sought to determine the decision-making model that would best explain estimated decision times and selection rate proportions. Evidence was based on the current correct probability for a target given the number of tokens within the left, right and center locations (**eq. 1**, see **Methods**). We tested five decision-making models (drift-diffusion model, drift diffusion model with leak, Trueblood (2021), urgency-gating model, urgency-gating model with a low-pass filter that used either novel evidence (**eq. 3**) or current evidence (**eq. 2**) to make a decision; see **Supplementary E**)^14,24^ Here we focus on **Experiment 3** (**Fig. 7**) since there was a significantly earlier estimated decision time in the slow rate token patterns compared to the fast rate token patterns (see **Supplementary E** for **Experiment 1** and **2** results).

**Figure. 7:**
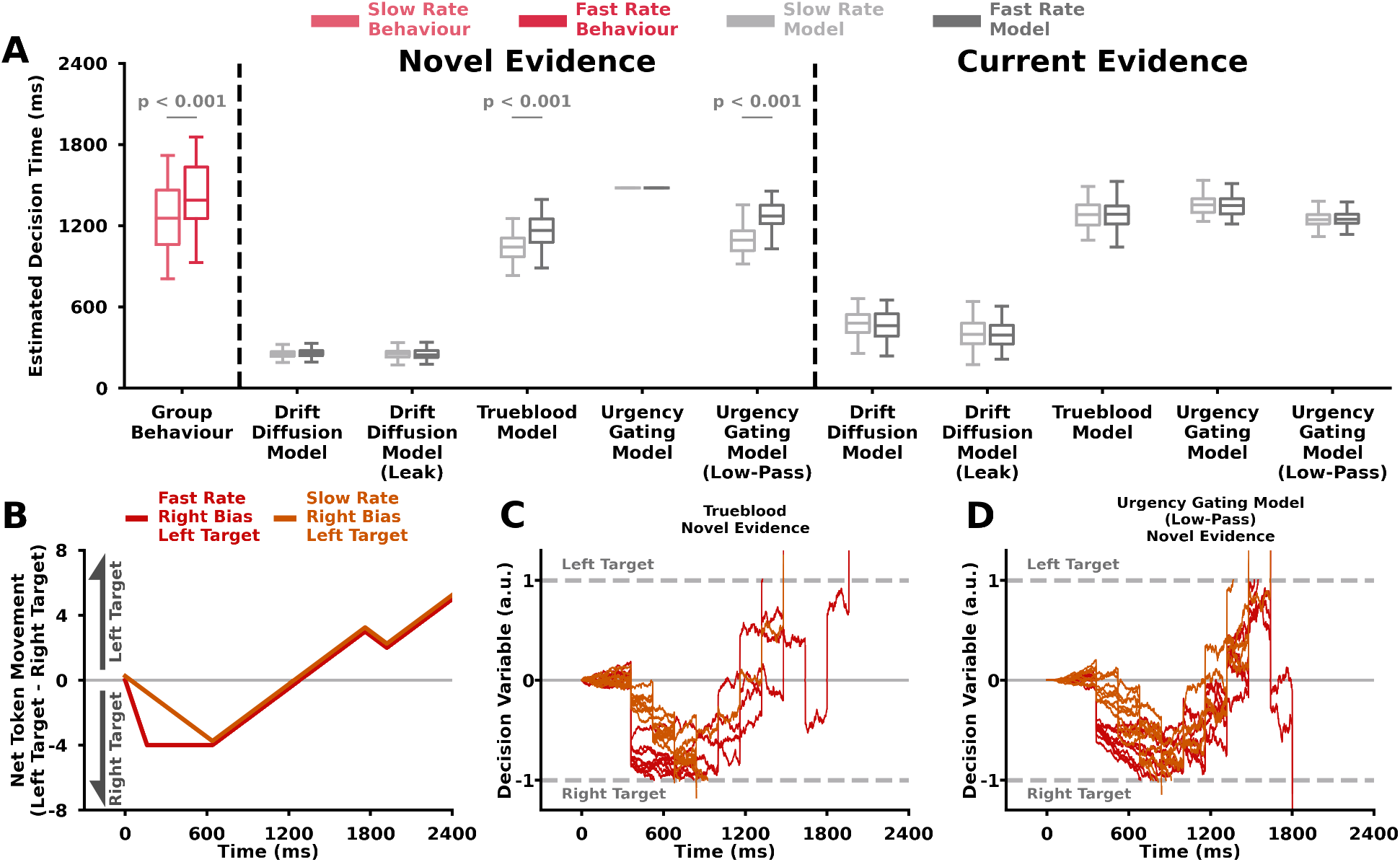
Best-Fit Decision-Making Model Simulations. **A) Experiment 3** estimated decision time (y-axis) for group behaviour (pink) and Decision-Making Models (grey; x-axis). Group participant estimated decision times are shown for slow rate token patterns (light pink) and fast rate token patterns (dark pink). Best-fit model simulations of decision times are shown for slow rate token patterns (light grey) and fast rate token patterns (dark grey). Box and whisker plots show 25%, 50% and 75% quartiles. Decision-making models simulated decisions using novel sensory evidence or current sensory evidence. As described above in **Figure 6**, participants made earlier decisions with slow rate token patterns compared to fast rate token patterns. Only the Trueblood model using novel sensory evidence and the urgency-gating model with a low-pass filter on novel sensory evidence were able to capture the behavioural difference in decision time between slow rate token patterns and fast rate token patterns. The Trueblood model and urgency-gating model with a low-pass filter both contain a temporally increasing (urgency) component and an integration of evidence. **B)** Net Token Movement (y-axis) over time (x-axis) for Slow Rate, Right Bias, Right Target (Dark Orange) and Fast Rate, Right Bias, Right Target (Dark Red) token patterns. **C-D)** Example simulations of decision-making models showing decision variables (y-axis) over time (x-axis). Each trace represents a single decision-making trial for either slow rate, right bias, right target (dark orange) and fast rate, right bias, right target (dark red) token patterns. The dashed grey lines represent decision thresholds for a left target decision or right target decision. **C)** Trueblood model using novel evidence. **D)** Urgency-gating model with a low-pass filter using novel evidence. Our model results suggest that the deliberation process likely includes an urgency signal, or temporal scaling, component as well as the integration of novel evidence.

We found the Trueblood model with novel evidence and the urgency-gating model with a low-pass filter with novel evidence were the only two models which could capture the earlier decision times in the slow rate token patterns relative to the slow rate patterns (**Fig. 7A**). The other best-fit models found decision times that were similar between the two different sets of rate token patterns.

To give insight into the mechanisms of the models, we show representative model behaviour in **Figure 7C-D**. In **Fig. 7B**, we show examples of fast rate right bias left target and slow rate right bias left target token patterns. These two token patterns were similar except for the different rates of token movement for the initial bias. For both the Trueblood model with novel evidence (**Fig. 7C**) and the urgency-gating model with a low-pass filter on novel evidence (**Fig. 7D**), we see similar decision variable trends. Both the Trueblood model and the urgency-gating model with a low-pass filter utilize urgency and integrate evidence leading to similar behaviour. For the fast rate token pattern there is some initial integration of evidence, either through evidence accumulation or the low-pass filter. However, urgency is low early when the first four tokens move, so that the decision variable does not immediately cross the decision threshold. Conversely for the slow rate token pattern, each individual token movement leads to some integration of evidence. Crucially, individual token movements later in time are more heavily weighted by urgency, which compounded over time leads to an earlier corsssing of the decision variable over the decision threshold. Note for the drift diffusion models, the best solution to capture the trend was achieved by having high noise parameters since they would be unable to produce the observed faster decision time with the slow rate token pattern. We chose to use the Trueblood model as an input into the decision-making and movement model, described directly below, because it explicitly defines both urgency and evidence accumulation.

#### Decision-Making and Movement Model

We found that ongoing deliberation influenced online movement. To capture this movement behaviour, we developed an optimal feedback control model^25,26,27,28,29,30^ that used the evolving decision variable to influence the ongoing movements. The decision variable was simulated using the Trueblood model with novel evidence. In short, an optimal feedback controller directed the hand towards an evolving and weighted averaged target that was a function of deliberation (see **Supplementary E** for further details). This model is able to capture individual movement behaviour (**Fig. 8A-C**) and group movement behaviour (compare **Fig. 8D-F** to **Fig. 8G-I**).

**Figure. 8:**
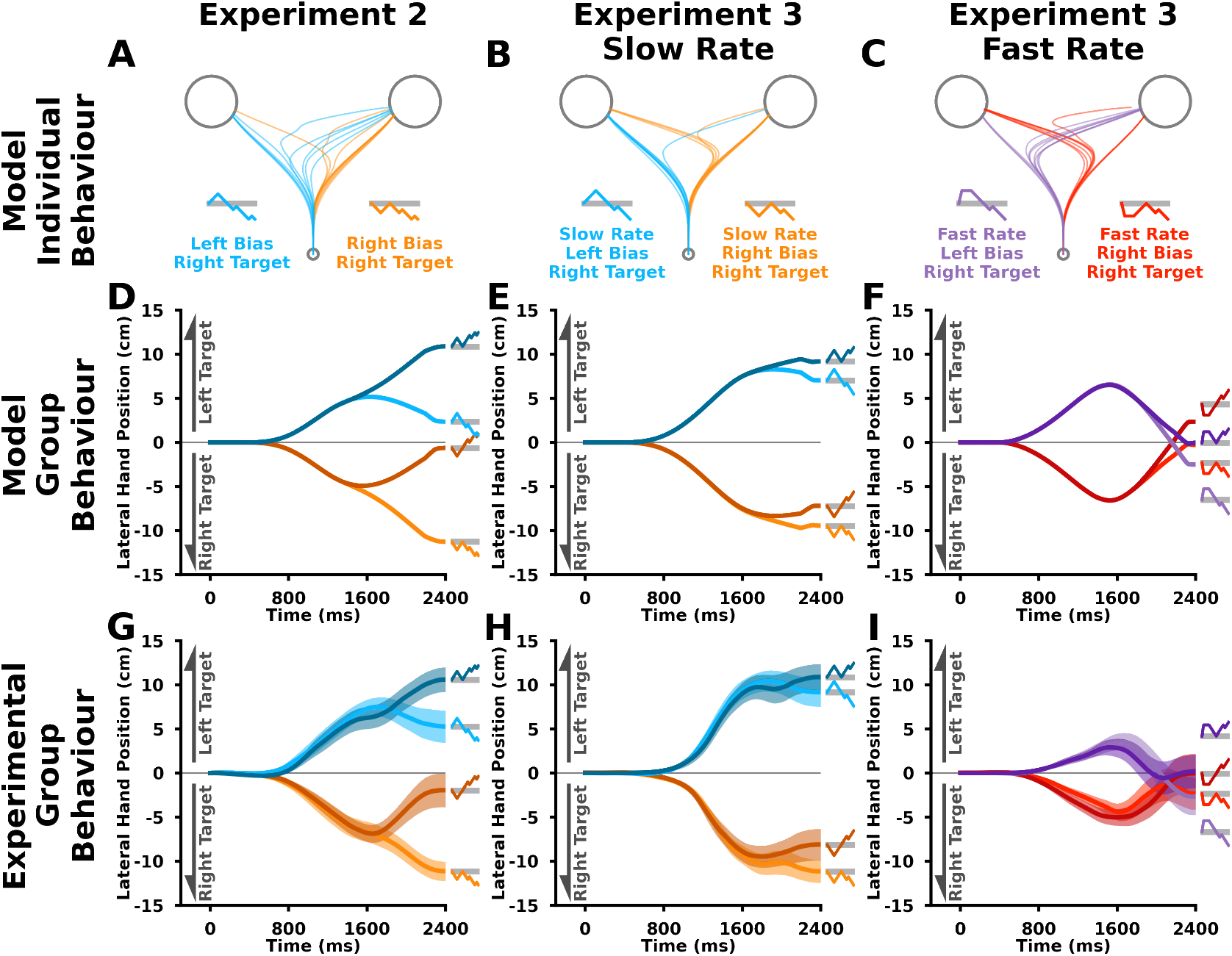
Best-Fit Movement Model Simulations. We fit a decision and movement model across the movement trajectories in biased token patterns in **Experiment 2** and **Experiment 3**. The models utilized a weighted average of the targets to control the feedback responses. For each trial and time step, the weighting for each target was calculated from a decision variable generated by the Trueblood model using novel sensory information. **A-C)** Model Individual Behaviour. **D-F)** Model Group Behaviour Lateral Hand Position (y-axis) over time (x-axis). **G-I)** Experimental Group Behaviour Lateral Hand Position (y-axis) over time (x-axis; repeated from Figure 4). The model was able to capture the trends found in the experimental group behaviour. This model supports the idea that online movements reflect the ongoing deliberation process.

#### Replicating previous work with the movement model

Using our decision-making and movement model, we were also able to replicate the results from a go-before-you-know task by Wong and Haith (2017).^3^ The researchers defined reaches that were not directly at one of the two targets as intermediate movements. They found that slow reaching movements resulted in more intermediate movements compared to fast reaches (**Fig. 9**). The authors interpreted these findings to indicate a single flexible plan that maximized task performance, since an averaging of static motor plans would always launch as an intermediate movement regardless of movement speed.^1^

**Figure. 9:**
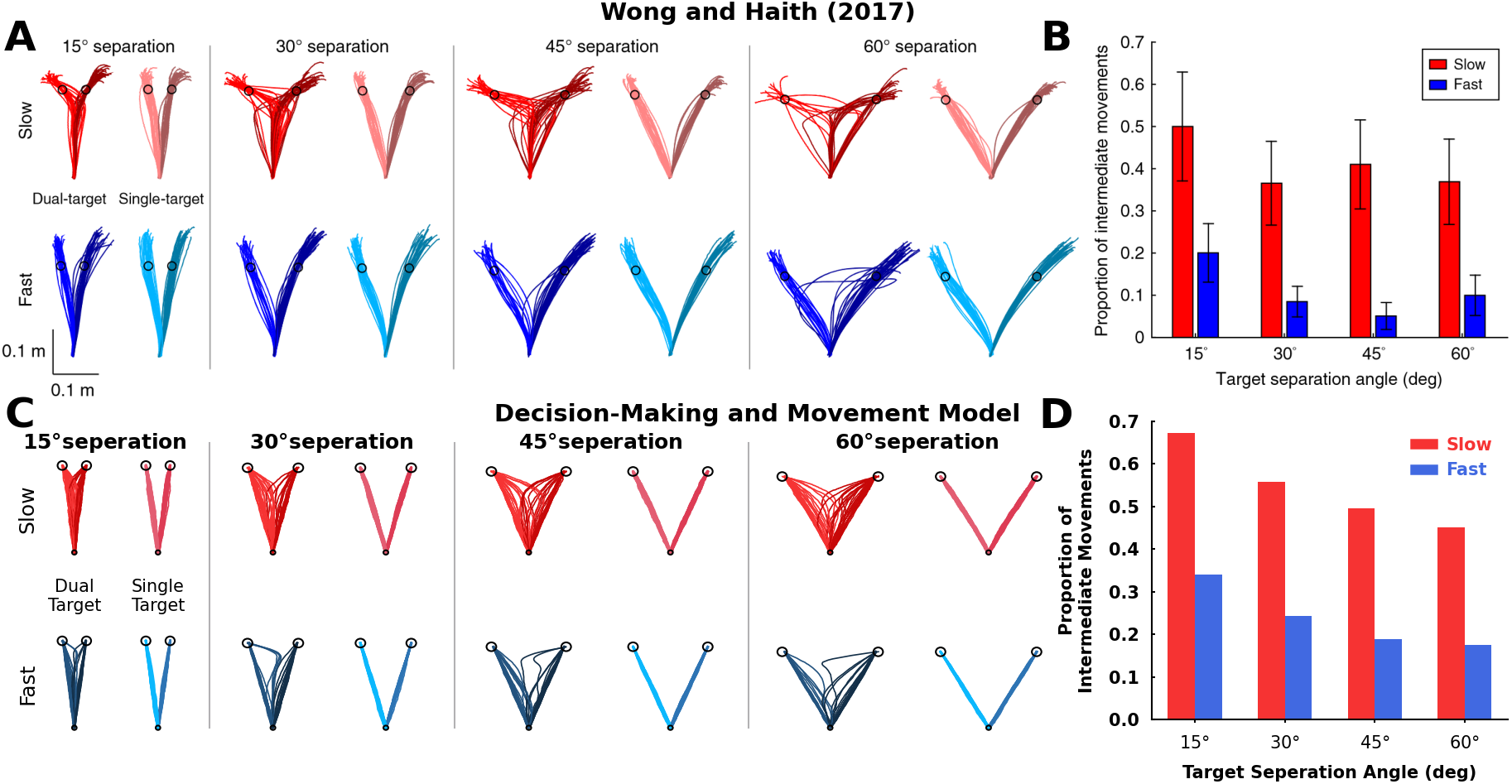
Replicating Previous Work. **A,B)** Behavioural data from Wong and Haith (2017; reprinted with permission) showing that required movement speed and target separation affect the proportion of intermediate movements. **A)** Individual single trajectories for dual-target and single target reaches with different target separation angles. Red and blue represent movements in the slow and fast conditions respectively. **B)** Group proportion of intermediate movements (y-axis) between target separation angles (x-axis). In **C,D)**, our model predicts that differences in urgency between conditions can lead to differences in the proportion of intermediate movements. Here, we modulated urgency as a function of the relative cost of an intermediate movement and the time to get to the target. **C)** Model single trajectories for dual-target and single target reaches with different target separation angles. Red and blue represent movements in the slow and fast conditions respectively. **D)** Model Average proportion of intermediate movements (y-axis) between target separation angles (x-axis). Our model was able to replicate the influence of required movement and target separation angle on the proportion of intermediate movements by manipulating the urgency in the deliberation process.

We replicated their findings (**Fig. 9C,D**) by using an urgency signal that was inversely proportional to allowable reach time, as well as proportional to the distance between the targets since this would be more energetically costly (**eq. 27**).^31^ In particular when comparing slow and fast movement speeds, our decision-making and movement model suggests that the proportion of intermediate movements arises due to the urgency to make a decision. For example, urgency is higher in the fast movement condition since there is less time to reach the target. As a consequence during these fast movements, a target is more quickly selected even without evidence, since the deliberation noise is multiplied by a high urgency signal and crosses a decision threshold (i.e., guessing). Conversely in the slow movement condition, the lower urgency does not push the noise over a decision threshold and the participant can wait for evidence of the correct target.

Collectively our empirical and computational results suggest that deliberation, which involves urgency, directly influences online movements.

## DISCUSSION

We show that ongoing deliberation is reflected in movements—prior to a decision—when the motor system is actively engaged. We also find that urgency was necessary to explain decision times in the third experiment, as well as predicting movement behaviour in the literature. Collectively, our work supports the idea that decision-making processes continuously interact with motor processes, such that deliberation is expressed via movement.

In **Experiment 2** and **Experiment 3**, we were able to elucidate the influence of the ongoing deliberation of uncertain and continuous evidence on movements. Prior literature has utilized a “go-before-you-know” paradigm where participants were presented multiple potential targets and initiated their movements without complete knowledge of the correct target.^7,1,32,33,2,3,4^ In these studies, the correct target was indicated partway through the reach via a sudden and discrete change of evidence (e.g. target colour or location, phonological input, etc.) that resulted in participants making rapid movement redirections. These rapid movement redirections reflect a rapid decision in response to a sudden and discrete change of evidence. Similar rapid movement redirections have been seen following uncertain and continuous evidence (i.e., random dot motion task) that is presented prior to movement initiation.^9,34,35^ In a small subset of trials, participants displayed “changes of mind” where they rapidly redirected towards the other target. It has been suggested that these changes of mind reflect a second decision based on delayed sensory information. Due to the sudden decisions and rapid movement redirections in the above works, it would be difficult to dissociate whether movement was caused either from deliberation or acting solely on a second decision.

There has also been increased interest in mid-reach decisions, such as when using the target-split paradigm by Kurtzer and colleagues (2020).^36^ In this task, participants would move their hand to one target and this would occasionally change to two target options during the movements. Participants showed a preference toward the options nearest the original target. Others have shown that mid-reach decisions are sensitive to other factors such as relative target frequency,^37^ reward magnitude,^38^ and biomechanics.^39,40,41^ In these mid-reach decision tasks, participants indicate their choice with a rapid movement redirection. Again however, it would be difficult to dissociate whether movement was caused either from deliberation or the final decision.

Unlike the above works and others,^42,43^ a key aspect of our design was using the hand trigger to estimate the decision time, allowing us to separate whether movement was caused either from deliberation or action selection following a final decision. Future work could adapt this paradigm during reaching or gait to study the influence of reward, energetics, and other factors that may impact decision-making to gain an understanding of the ongoing deliberation via movement.

In **Experiment 1** we found that the ongoing deliberation did not induce movements prior to a decision, at least to a significant level, when initially in posture. Conversely, in both **Experiment 2** and **Experiment 3** we found that when the motor system was already actively engaged that there was an expression of the deliberation process via movement. Being able to express deliberation in movement but not in posture aligns with previous results showing differential configuration and engagement of motor circuits for movement and posture.^44,45,46^ One possibility is the decision processes have a larger influence on movement circuits than postural circuits. While our paradigm allows for a continuous expression of deliberation during movement execution, past work has shown that it is possible to elicit an instantaneous expression of deliberation from a postural state. Selen and colleagues were able to gain a momentary expression of deliberation at the moment of movement onset.^47,34^ Specifically, they perturbed the upper limb while in posture and measured the resulting long-latency stretch reflex. They found that the long-latency stretch reflex reflected deliberation at the time of perturbation while in posture. Although we did not find that evidence was enough to elicit movement initiation from a postural state, we did find that deliberation can be continually expressed during movement.

In this work, we have primarily investigated the influence of deliberation on movement. We also found that participants made faster decisions when already moving in **Experiment 2** compared to when in posture for **Experiment 1**. This finding may reflect ‘embodied decisions, where the current and future states of the motor system can influence decision making.^48,49,50,51,52,39,53,54,55,56,41^ Korbisch and colleagues (2022) had participants select between short or long walking durations or shallow and steep walking inclines.^54^ When participants looked at depictions of the various options, the researchers found higher saccade vigor (i.e., velocity) towards the depictions associated with less effort. These results suggest that potential energy costs are embodied and can be reflected during deliberation with eye movement. In their study, evidence of potential effortful options was discrete and did not change during the course of the eye movement. Here saccade vigor provides a glimpse of deliberation reflecting past evidence acquired from previous eye movements. Building on this work, we show the online movement itself is influenced by an ongoing deliberation. It would be interesting for future work to manipulate both potential energetic costs over time and evidence during movement to further understand embodied decisions.

In this study, we were interested in the influence of deliberation on movement. In **Experiment 1** and **Experiment 2** we found no difference in decision times between the bias token patterns, which replicates previous findings and is consistent with the urgency-gating hypothesis..^14^ For **Experiment 3**, we used slow rate and fast rate token patterns to manipulate the rate of evidence and further understand deliberation. The standard evidence accumulation (with or without leak) and pure urgency-gating model (without a low pass filter) would predict that the fast rate token patterns would respectively cause earlier or similar decision times compared to the slow rate token patterns. Counterintuitively, we found that the slow rate token patterns made faster decisions compared to the faster rate token patterns. We were able to capture faster decisions with the slow rate token patterns with both the Trueblood model and urgency-gating model with a low-pass filter. Both these models are similar mathematically and have terms that relate to urgency and an integration of evidence. Conceptually, the Trueblood model integrates to accumulate evidence towards a decision, whereas the integration from the low-pass filter of the urgency-gating model is intended to reflect an estimate of evidence from sensory processes. Neural activity during perceptual decision-making in monkeys has been attributed to either evidence accumulation towards a decision.^17,11,21^ or the scaling of low-pass filtered estimate of evidence with an urgency signal that arises from the basal ganglia.^14,16^ An important future direction, such as through neural recordings in animals, is to determine where and why there is an integration of evidence. Irrespective of evidence integration, urgency was needed to predict decision times and replicate reaching trajectories from past work.^3^

Here developed a movement model that reflected deliberation, by combining the Trueblood decision-making model and optimal feedback control. This differs from past work that has used dynamic programming,^57^ bayesian methods,^58^ only optimal feedback control,^49^ and relative desirability of multiple options.^59^ While these other modelling approaches have been insightful and motivated the current work, they do not have a deliberation process that includes urgency. Urgency was a critical component to capture decision making time and reaching movements from past literature. However, it would be possible to include an urgency term in these previous modelling approaches. A limitation of our model as currently formulated is that it only allows for the deliberation process to influence the movement. That is, it does not allow the motor states to directly influence the deliberation process. This model design reflects our experiments where we manipulate the deliberation process to test its influence on movement. However, several of the aforementioned models would be able to capture some of the bidirectional relationships between cognitive and motor processes during embodied decisions reported in the literature.^48,51,53,52,54,41^ Moving forward, it will be important to have a computational model of embodied decisions that captures several important features of both motor behaviour (e.g., bell shaped velocity profiles, vigor) and decision-making behaviour (e.g., skewed reaction time, speed accuracy tradeoff, Hicks law, urgency).

Overall, we have shown that the motor system is influenced by the deliberation of multiple targets. Prior literature has examined how the decision-making and motor systems interpret and act on multiple potential options.^4,3,60,6^ In the go-before-you-know task, intermediate movements between two targets have been suggested to be an outcome of parallel averaged motor plans^1,2,59^ or a single flexible motor plan that optimizes task performance.^3,61,4^ Wong and Haith (2017) interpreted more intermediate movements with slow hand speeds compared to fast hand speeds to reflect a single flexible motor plan.^3^ Here we provide an alternative perspective by considering urgency. When one also considers urgency, it is possible to explain different proportions of intermediate movements between slow or fast hand speeds with either a single flexible motor plan or parallel averaged motor plans.

It is important to consider that a single flexible motor plan or parallel averaged motor plans are a combination of two factors: i) single versus parallel average, and ii) static versus flexible. Obviously a single static motor plan is not a viable option to handle multiple potential goals. Alhussein and colleagues (2021) rule out a parallel average of static motor plans, since their prediction was based on the initial reach angle to each target.^4^ Yet their finding does not rule out the possibility of a parallel average of flexible motor plans, where each motor plan (more specifically, control policy) could contain a safety margin. As shown above, we were able to replicate the results of Wong and Haith (2017) by considering urgency.^3^ It is mathematically equivalent to have a single flexible motor plan that reflects a weighted average of two targets based on evidence, compared to flexible parallel plans (control policies) that are weighted based on evidence (see **Supplementary E**). It is not clear how to behaviourally dissociate between a single flexible motor plan or parallel average of flexible motor plans through movement execution. There has been conflicting neural support with regards to parallel motor plans or a single flexible motor plan.^5,6^ It would be useful for future work involving neural recordings to determine where, when, and how multiple target representations and deliberation processes finally converge to produce a single executed movement.

Humans often must make decisions while moving. We found that deliberation was reflected in ongoing movements—prior to a decision—when the motor system was actively engaged. We found that an urgency signal, which more heavily weighted evidence later in time, was fundamental to predicting decision times and explaining previous reaching behaviour. Our results support the hypothesis that the decision-making process influences movements prior to a decision. Understanding the integration of decision and motor processes may allow us to better understand neurological disorders where cognitive and motor processes and deficits may be entangled.

## METHODS

### Participants

In total we collected 51 participants across three experiments. 17 individuals (24.8 ± 2.37 years old) participated in **Experiment 1**, 17 individuals (21.4 ± 1.76 years old) participated in **Experiment 2**, and 17 individuals (23.2 ± 2.93 years old) participated in **Experiment 3**. Participants reported they were free of musculoskeletal or neuromuscular disorders. All participants provided informed consent to participate in the experiment and the procedures were approved by the University of Delaware’s institutional review board. Participants were provided $10 USD compensation.

### Apparatus

For all three experiments, participants grasped the handle of a robotic manipulandum with their dominant hand (**Fig. 1A**; KINARM, BKIN Technologies, Kingston, ON, Canada) to perform reaching movements in the horizontal plane. Participants held a hand trigger in their nondominant hand. A semi-silvered mirror projected images (start position, left and right targets, tokens) from an LCD screen onto the horizontal plane of motion. To assess muscle activity, we recorded electromyography (EMG) signals with bipolar surface electrodes (single differential electrode, Trigno system, Delsys, Natick, MA) from the flexor policis brevis of the nondominant hand. To obtain an estimated decision time, a voltage signal indicated when the thumb pushed the hand trigger. Kinematic, EMG, and hand trigger data were recorded at 1,000 Hz and stored offline for data analysis.

### Protocol

#### General Task Protocol

For each trial, participants were visually presented with a white start position (2 cm diameter) and two targets (5 cm diameter). The left and right targets were respectively 20 cm to the left and right of the start position (**Fig. 1A**). A yellow cursor (1 cm diameter) provided real-time feedback of their hand position. The participants were instructed to move their cursor into the start position. After holding the cursor with the start position for 400 ms, participants heard a beeping sound and 15 yellow tokens appeared between the left and right targets. At trial onset (0 ms), the tokens jumped from the center into the left target or right target in 160 ms time intervals^14^ (**Fig. 1C**). Participants had to make their decision prior to 2400 ms, corresponding to the final token moving into one of the targets. Once they felt confident which target would end up with the most tokens, they were instructed to simultaneously i) press a trigger with their non-dominant hand and ii) move towards and hit the selected target. As soon as participants pressed the hand trigger, the remaining token movements were not visible to the participant to prevent them from changing their decision with later evidence. If participants selected the correct target, they would hear a pleasant ding and their selected target would turn blue. If participants selected the incorrect target, they would hear an unpleasant buzzer and their selected target would turn red. When participants did not press the hand trigger and/or enter a target within 2400 ms of the beginning of the trial, both targets would turn red. Further, unknown to participants, that the trial would be repeated later on during the experiment.

#### Experiment 1 Task Protocol

The goal of **Experiment 1** was to determine if ongoing deliberation can elicit and subsequently influence movements, prior to a final decision, when evidence was initiated during posture. The targets were directly to the left and right of the start position (**Fig. 1A**). The participant waited in the start position for 400 ms. After this wait period, trial onset (0 ms) was indicated with a beep. The tokens moved into the left or right target one at a time in 160 ms intervals. In total, participants experienced 216 trials in the main experiment. We used bias, pseudorandom, late, and null token patterns (**Fig. SA1**).

We were primarily interested in the bias token patterns, since we tightly controlled the token movement and consequently the experienced uncertain and continuous evidence. During the bias token patterns the first three tokens moved individually into the left or right target (i.e., left bias or right bias), the next three tokens moved individually into the opposite target, and the remaining tokens moved with an 80% probability into the left or right target (i.e., left target or right target; **Fig. 2A-D**). These bias token patterns, we had each of the four combinations of left bias or right bias and left target or right target. Each bias token pattern was presented 12 times, which resulted in 48 bias token patterns.

We also had psuedorandom token patterns where each token had the same probability of going to the left target. We had 20%, 35%, 50%, 65% or 80% probability psuedorandom token patterns. Each psuedorandom token pattern was presented 12 times except for the 50% condition which was presented 24 times, which resulted in 72 psuedorandom token patterns. Additionally, we had null token patterns (24 trials), late token patterns (48 trials), and late null token patterns (24 trials). Similar to the ambiguous token patterns used by Cisek (2009), the null bias token patterns had a net token movement that was close to zero throughout the beginning portion of a trial.^14^

#### Experiment 2 Task Protocol

The goal of **Experiment 2** was to determine if ongoing deliberation was reflected in movements, prior to a final decision, after movement onset when the motor system was already actively engaged. Tokens were initiated when the participant left the when evidence was initiated by movement. In **Experiment 2**, the targets were 30 cm forward and 20 cm to the left and right of the start position. The participant waited in the start position for 400 ms, after which they heard a beep. The beep indicated the participant may leave the start position. Trial onset (0 ms) occurred once the participant left the start position. **Experiment 2** used the same token patterns as **Experiment 1**.

#### Experiment 3 Task Protocol

The goal of **Experiment 3** was to replicate the results found in **Experiment 2**, while also elucidating the roles of evidence accumulation or urgency on deliberation and consequent movement. The experimental setup was the same as **Experiment 2**, except for the specific token patterns (**Fig. SA 2**). Participants experienced 336 total trials. Trials included slow rate bias (**Fig. 2 E-F**), fast rate bias (**Fig. 2I-L**), pseudorandom, late, and null token patterns.

In this experiment, we were primarily interested in the slow rate and fast rate bias token patterns because we tightly controlled their movement and the experienced uncertain and continuous evidence. Further, the slow rate and fast rate token patterns lead to unique decision times depending on how humans accumulate evidence and / or rely on urgency during deliberation. During the slow rate bias token patterns, the first four tokens moved individually into the left or right target (i.e., left bias or right bias), the next four tokens moved individually into the opposite target and the remaining tokens moved with an 80% probability into the left or right target (i.e., left target or right target; **Fig. 2E-F**). In the fast rate bias token patterns the first four tokens moved at the same time into the left or right target (i.e., left bias or right bias), the next four tokens moved individually into the opposite target and the remaining tokens moved with an 80% probability into the left or right target (i.e., left target or right target; **Fig. 2I-L**). For these bias token patterns, we had each of the eight combinations of fast rate or slow rate, left bias or right bias, and left target or right target. Each bias token pattern was presented 12 times, which resulted in 96 bias token patterns.

The pseudorandom token patterns were the same as **Experiment 1** and **2** (**Fig. SA 2I-M**). Similar to **Experiment 1** and **2**, we also had late and null token patterns.

#### Reaction Time Task

Prior to any of the experiments described above, participants performed a reaction time task to determine the sensory and motor delays involved in making and indicating a decision (**Fig. SA 3A**). In the reaction time task, the targets were in the same location as the corresponding main experiment (as described in **Experiment Task Protocols** above). The reaction time task used the same trial onset as the corresponding experiment. At trial onset (0 ms), all 15 tokens jumped into either the left or right target (**Fig. SA 3B,C**). Participants were instructed to select the target that all of the tokens jumped into as fast as they could (**Fig. SA 3D**). Again, participants indicated their decision by pressing the hand trigger and moving the cursor into their selected target (**Fig. SA 3E,F**). Participants performed at minimum 20 familiarization trials in the reaction time paradigm to become accustomed to the experimental setup. After the familiarization trials, participants performed 24 reaction time trials. There were 12 left reaction time trials and 12 right reaction time trials that were presented in a randomly interleaved order.

### Data Analysis

#### Estimated Decision Time

Trigger time was determined when the voltage of the hand trigger crossed 3 volts for each trial. We found an estimated decision time on each trial to determine when decisions were made independent of reaching movements. We estimated a Neural + Mechanical Delay for each participant using their reaction time trials. For each muscle per trial, we subtracted the global mean muscle activity across all the reaction time trials. Flexor policis brevis muscle activity was full wave rectified and then dual-pass, sixth order, lowpass (20 Hz), Butterworth filtered. We determined EMG onset time with a dual-threshold method given a critical amplitude threshold and a 10 ms temporal threshold.^62^ We defined a critical amplitude threshold of mean + three standard deviations of the flexor policis brevis muscle activity in the 400 ms before the trial onset across all trials. EMG onset time was determined when the EMG activity rose and stayed above the critical amplitude threshold for 10 ms. The onset time was calculated using the dual-threshold method and verified by human inspection per reaction time trial (**Fig. SA 4A,B**). We found the average difference between Trigger Time and EMG onset time for the reaction time trials per subject (**Fig. SA 4C**). The Neural + Mechanical delay for each participant was defined as the average difference between Trigger Time and EMG onset time plus a nerve propagation delay of 20 ms.^63^ We calculated the estimated decision time on each trial during the main experiments as the trigger time minus the neural + mechanical delay (**Fig. SA 4D**).

#### Movement Analysis

Hand position data were digitally dual-pass, second order, lowpass (20 Hz cutoff), Butterworth filtered. Our primary focus was to determine whether the deliberation process influences movements, prior to a final decision. We were interested in the movement prior to the influence of the final decision and subsequent actions. To this end, we looked at the lateral hand position at estimated decision time (**Fig. 2**).

### Statistical Analysis

All statistical tests were performed in Python 3.8.5. We used repeated measures analysis of variance (rmANOVA) as the omnibus tests for each dependent variable. We were primarily interested in estimated decision time, lateral hand position at estimated decision time, and selection rate metrics for the bias token patterns. In **Experiment 1** and **Experiment 2**, we used a 2 (Bias: Left or Right) x 2 (Target: Left or Right) rmANOVA for decision time, lateral hand position at estimated decision time, and selection rate. In **Experiment 3**, we used a 2 (Rate: Fast or Slow) x 2 (Bias: Left or Right) x 2 (Target: Left or Right) rmANOVA for decision time and selection rate. For lateral hand position at estimated decision time we performed separate 2 (Bias: Left or Right) x 2 (Target: Left or Right) rmANOVAs for fast bias patterns and slow bias patterns. Here we used separate rmANOVAs, since we found significantly different decision times between slow rate and fast rate bias token patterns. For **Experiments 1, 2**, and **3**, we were also interested in the pseudorandom token patterns and used a 1-way rmANOVA (Probability of Left Target: 20%, 35%, 50%, 65%, and 80%) for estimated decision time, lateral hand position at estimated decision time, and selection rate. For all experiments, we performed nonparametric bootstrap hypothesis testing for mean comparisons (n = 1,000,000).^64,65,66,67,68^ Holm-Bonferroni corrections were used to control for Type 1 error. We computed Common Language Effect Size 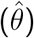 for all mean comparisons.^69,68^ Statistical significance was set to p < 0.05.

## Supplementary A

### Methods

#### Token Patterns

**Figure SA1:**
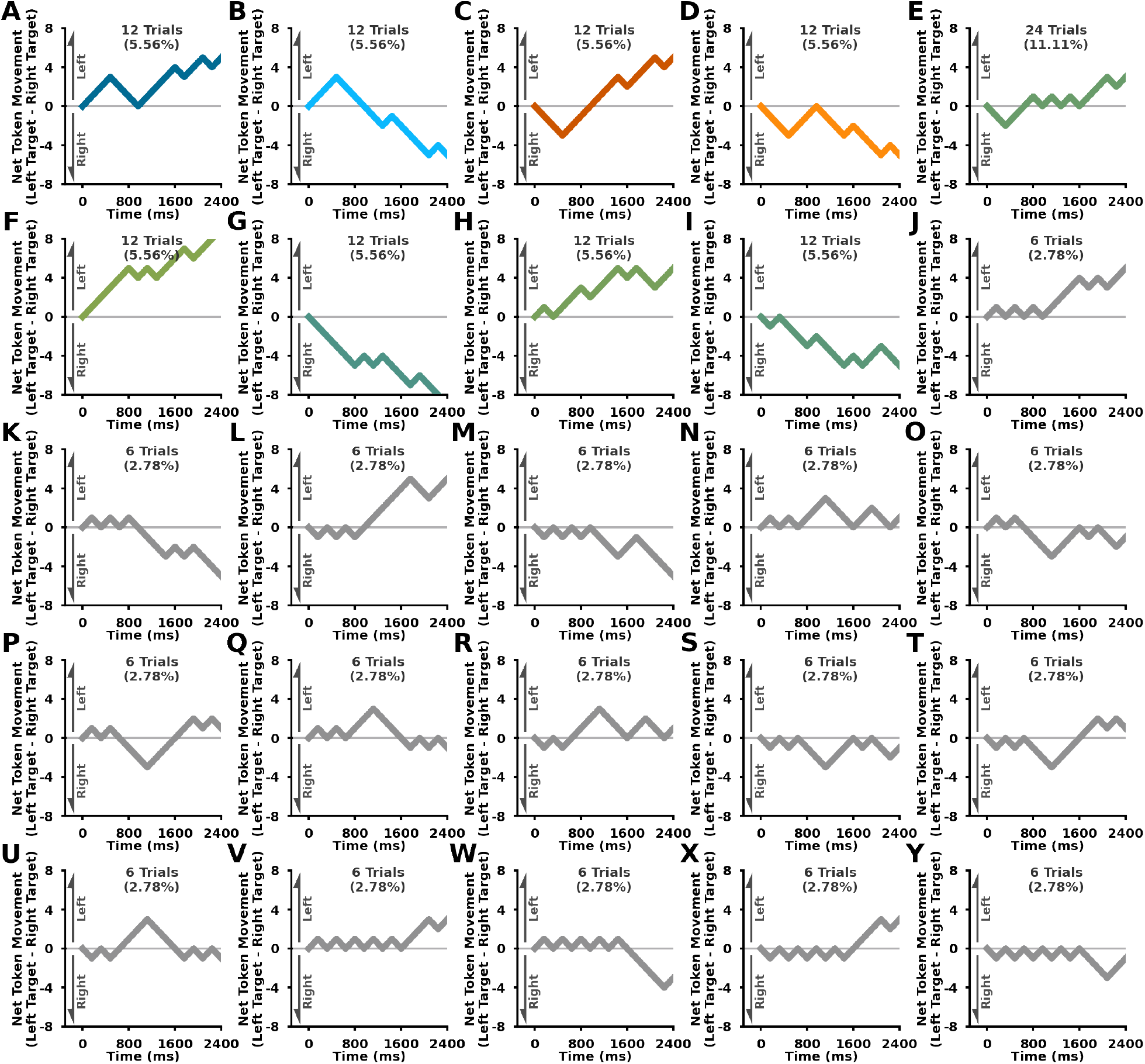
Experiment 1 and 2 Token Patterns. Net Token Movement (y-axis; left target - right target) over time (x-axis). Inset text for each token pattern shows the number of occurrences and percentage out of all trials.

**Figure SA2:**
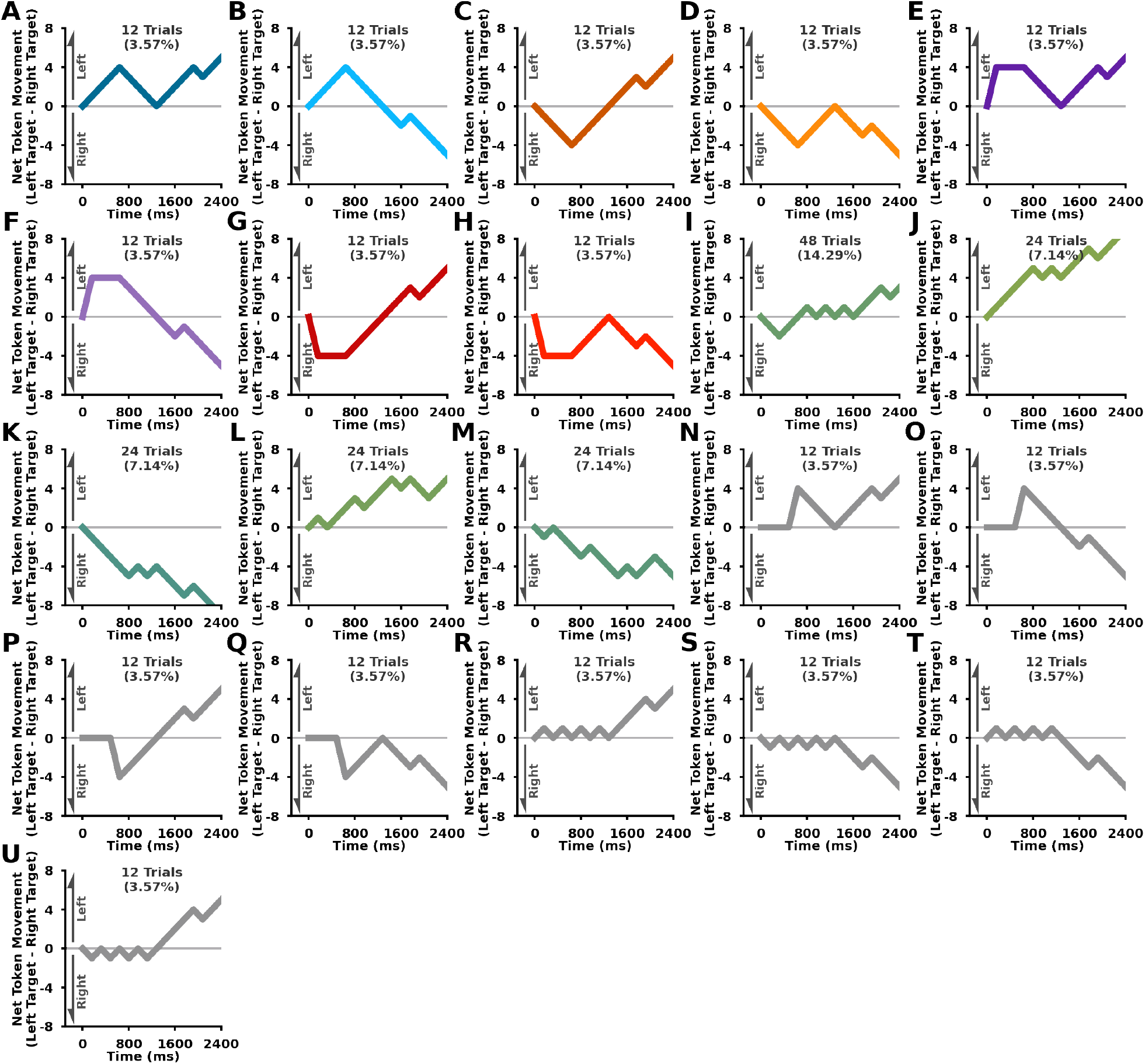
Experiment 3 Token Patterns. Net Token Movement (y-axis; left target - right target) over time (x-axis). Inset text for each token pattern shows the number of occurrences and percentage out of all trials.

#### Reaction Time Task

**Figure SA3:**
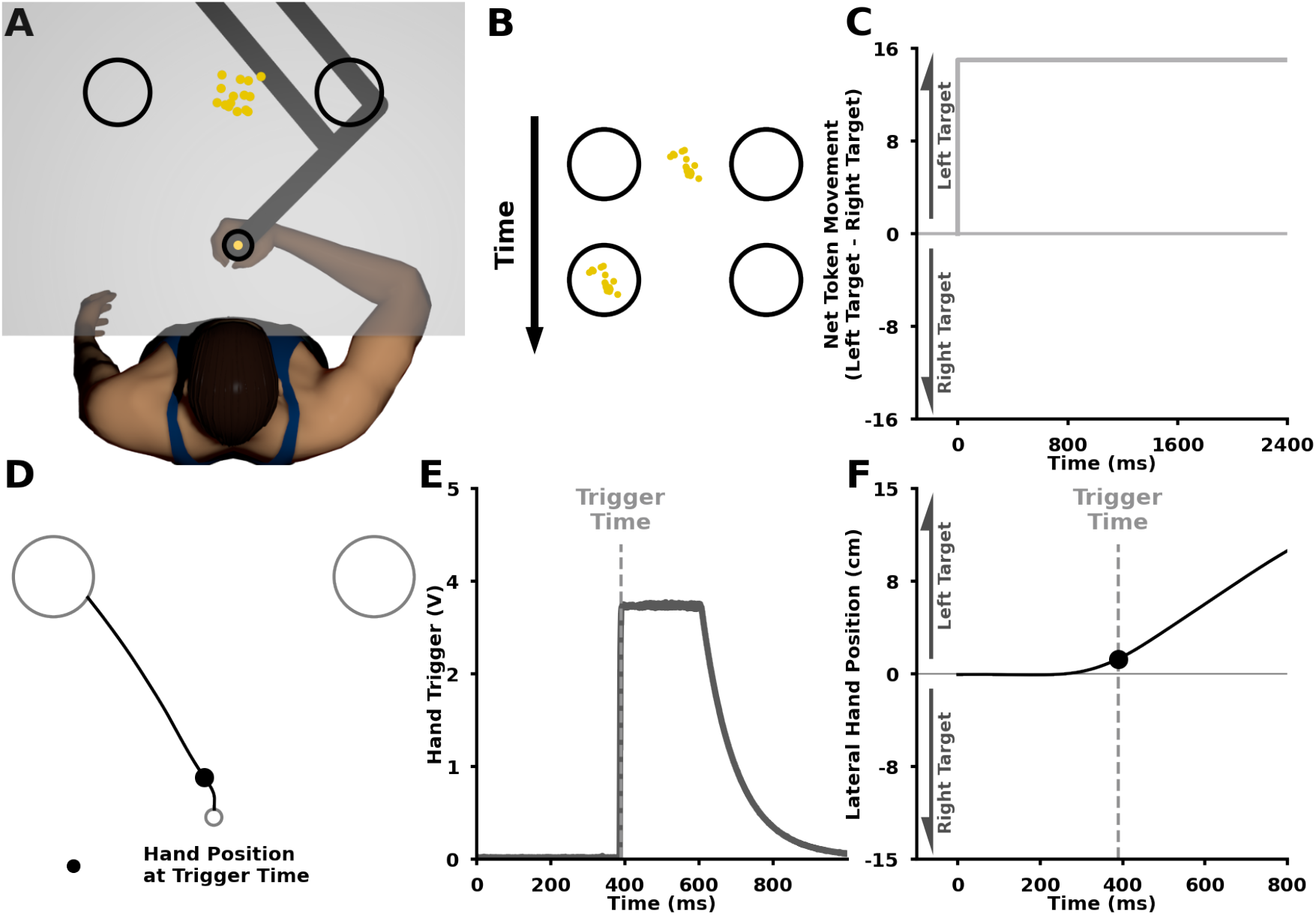
Reaction Time Design. In the reaction time trials, we wanted to measure how quickly participants could respond to goal-related stimulus. The participant initiated the reaction time trial by leaving the start position. As soon as participants left the start position, all the tokens jumped into one of the two targets. Participants were instructed to select the target which all of the tokens moved into as fast as they could. The participants indicated their by pressing the hand trigger in their non-dominant hand and moving into the corresponding target. **A)** The reaction time task setup was identical to the experimental conditions for **Experiment 2** and **Experiment 3. B)** An example of the participant display while the tokens moved into the left or right target over time. **C)** Net Token Movement (left tokens minus right tokens, y-axis) over time (x-axis) of an example token pattern. **D)** Individual reaction time trial reaching trajectory. Solid black circle represents the hand position at the trigger time. **E)** Hand trigger voltage (y-axis) over time (x-axis) for the trial shown in (**D**). The trigger time was the defined as the first time point the hand trigger voltage crossed a 3V threshold. **F)** Lateral hand position (y-axis) over time (x-axis) for the trial shown in (**D**). The vertical grey line in (**E**) and (**F**) indicates the measured trigger time.

#### Estimated Decision Time

**Figure SA4:**
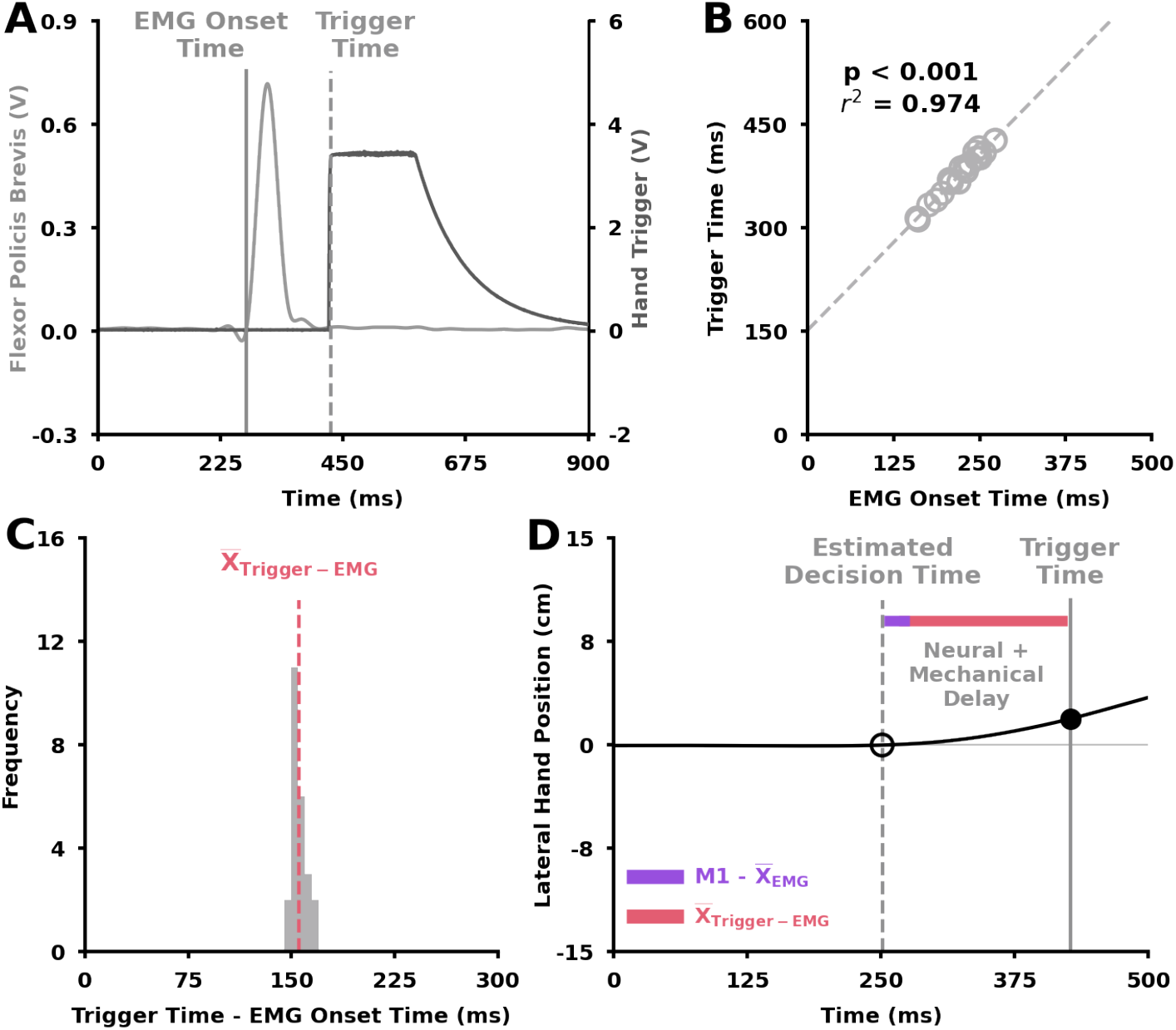
Neural and Mechanical delay calculation for example participant. **A)** Example single reaction time trial behaviour. Flexor policis brevis activity (left y-axis; light grey) and hand trigger voltage (right y-axis; dark grey) over time (x-axis). Vertical solid grey line represents EMG onset time. Vertical dashed grey line represents trigger time. **B)** Single participant trigger time (y-axis) vs EMG onset time (x-axis) for all reaction time trials. **C)** Histogram of difference between trigger time and EMG onset time (x-axis) for example participant reaction time trials. Vertical dashed pink line represents average difference between trigger time and EMG onset time. Average difference between trigger time and EMG onset time is used to calculate estimated decision time for each trial in experimental conditions. **D)** Lateral hand position (y-axis) over time (x-axis) for example reaction time trial. We define the neural + mechanical delay as the sum of average difference between trigger time and EMG onset time and an estimated 20ms neural propogation delay from the M1 brain region to the flexor policis brevis. From the trigger time on a single trial, we subtract the neural + and mechanical delay to calculate the estimated decision time. The estimated decision time allows us to look at the influence of ongoing deliberation prior to final decision-related movements.

## Supplementary B

### Conservative Estimate

**Figure SB1:**
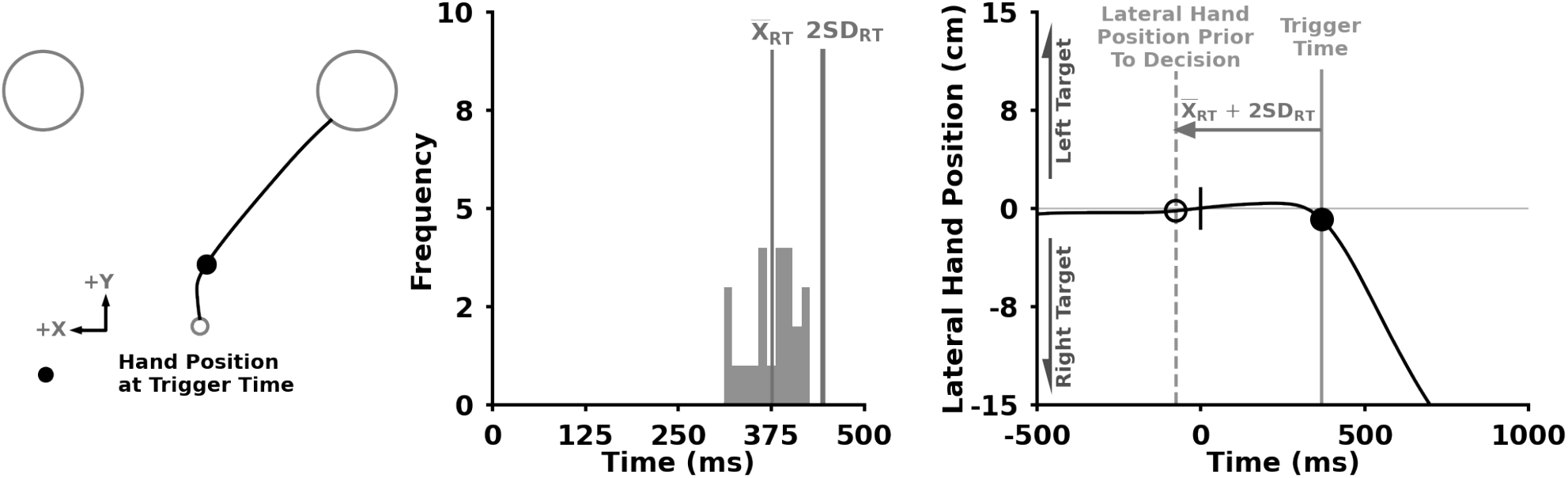
Conservative Time Estimate Prior to a Decision Example. **A)** Example single trial trajectory for a reaction time task. **B)** Histogram of trigger time for reaction time trials for a single participant. Vertical lines represent mean reaction time and mean + 2 standard deviations of reaction time. The mean + 2 standard deviations of reaction time are used to conservatively estimate a time point prior to any decision related behaviour. **C)** Lateral hand position (y-axis) over time (x-axis) for example reaction time trial. 0 ms is when participants left the start position in the reaction time trial. Vertical solid grey line represents the trigger time. Vertical dashed grey line represents the conservative time estimate prior to a decision. The short black line at 0 ms represents the beginning of the reaction time trial. We subtract the mean + 2 standard deviations of reaction time from the trigger time to conservatively estimate a time point prior to any final decision-related behaviour. In this trial, our conservative estimate is prior to the beginning of the reaction time trial.

### Lateral Hand Position Prior to Decision

**Figure SB2:**
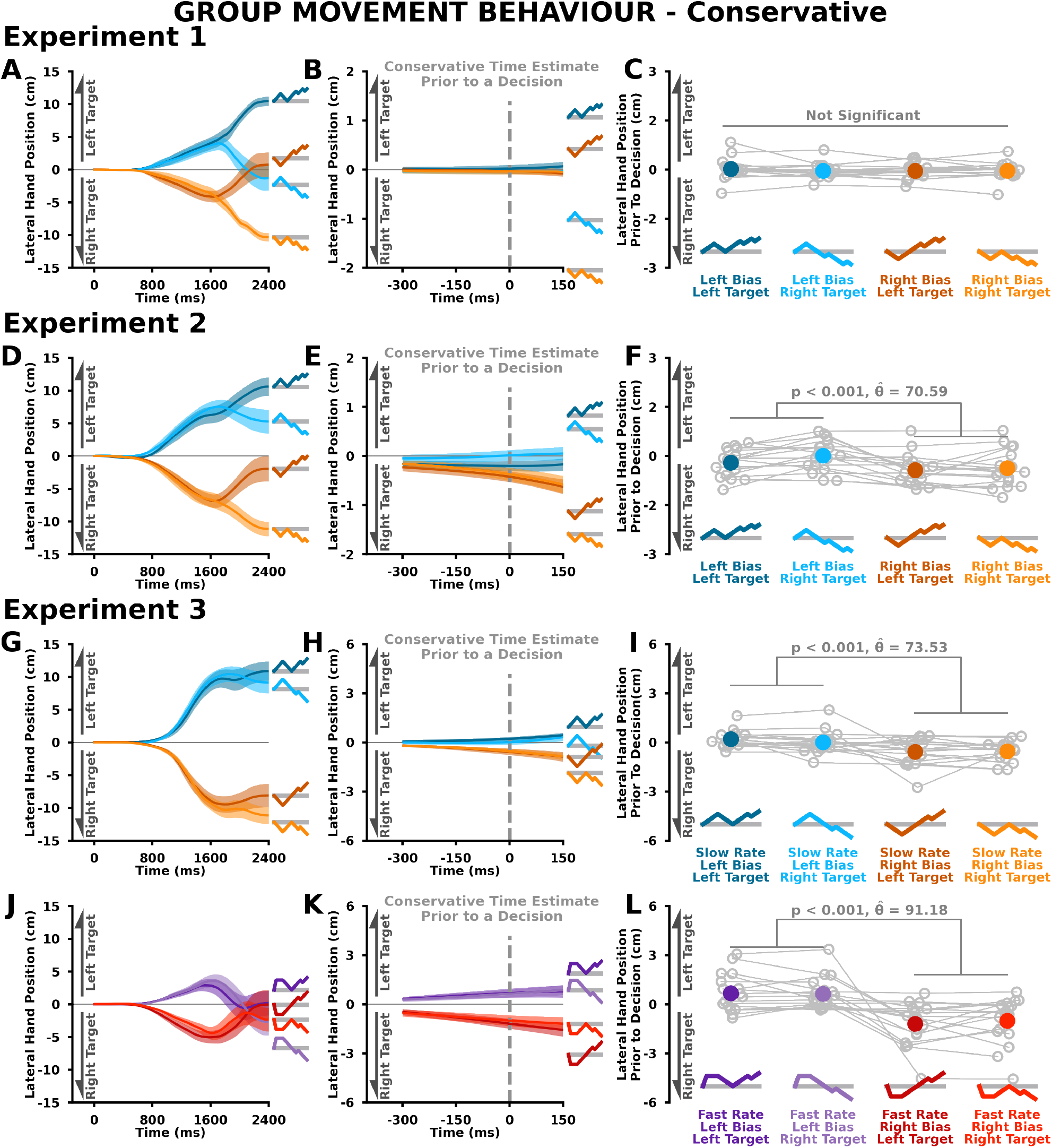
Group Movement Behaviour relative to Conservative Time Estimate Prior to a Decision. The figure is similar to **Figure 5** but for movement behaviour at the conservative time estimate prior to a decision. The results for the conservative time estimate prior to a decision are consistent with the results found using the estimated decision time (**Figure 5**).

## Supplementary C

### Group Behaviour - Psuedorandom Token Patterns

#### Group Movement Behaviour

**Figure SC2:**
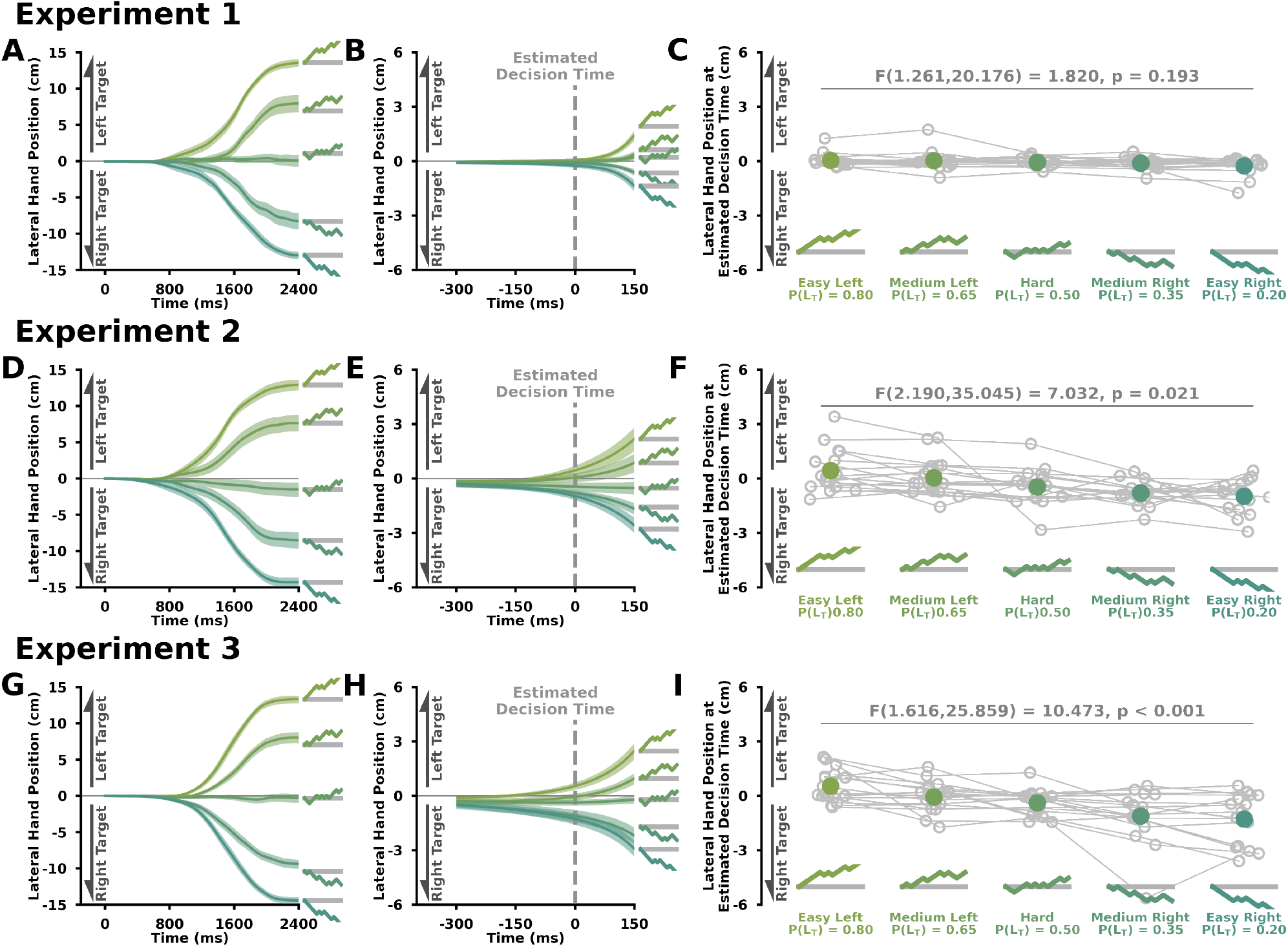
Group Movement Behaviour for Pseudorandom Token Patterns. **A,D,G)** Lateral hand position (y-axis) over time (x-axis) for pseudorandom token patterns in **A) Experiment 1, D) Experiment 2**, and **G) Experiment 3. B,E,H)** Lateral hand position (y-axis) over time (x-axis) aligned to estimated decision time for pseudorandom token patterns in **B) Experiment 1, E) Experiment 2**, and **H) Experiment 3. C,F,I)** Lateral hand position (y-axis) at estimated decision time across pseudorandom token patterns (x-axis) in **C) Experiment 1, F) Experiment 2**, and **I) Experiment 3**. Inset text shows the f-statistic for a main effect of the pseudorandom token pattern from an rmANOVA. These results are consistent with the findings shown in **Figure 4**. Again we see an influence of the token patterns on the movement prior to a decision in **Experiment 2** and **Experiment 3** but not **Experiment 1**.

#### Decision Time

**Figure SC1:**
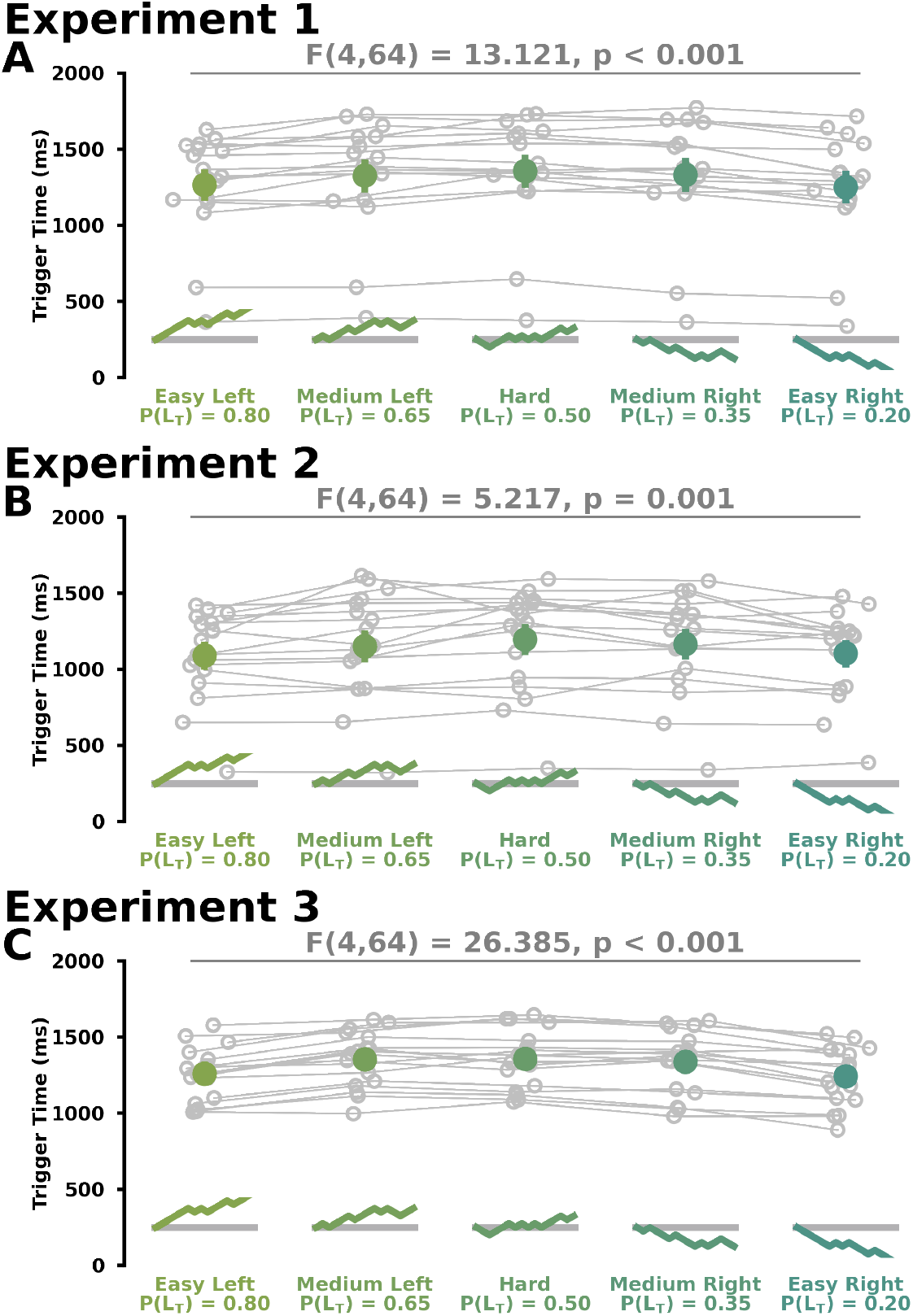
Group Estimated Decision Time Behaviour for Pseudorandom Token Patterns. Estimated decision time (y-axis) across pseudorandom token patterns (x-axis) in **A) Experiment 1, B) Experiment 2**, and **C) Experiment 3**. Inset text shows significant effects from an rmANOVA.

#### Group Selection Rate Behaviour

**Figure SC3:**
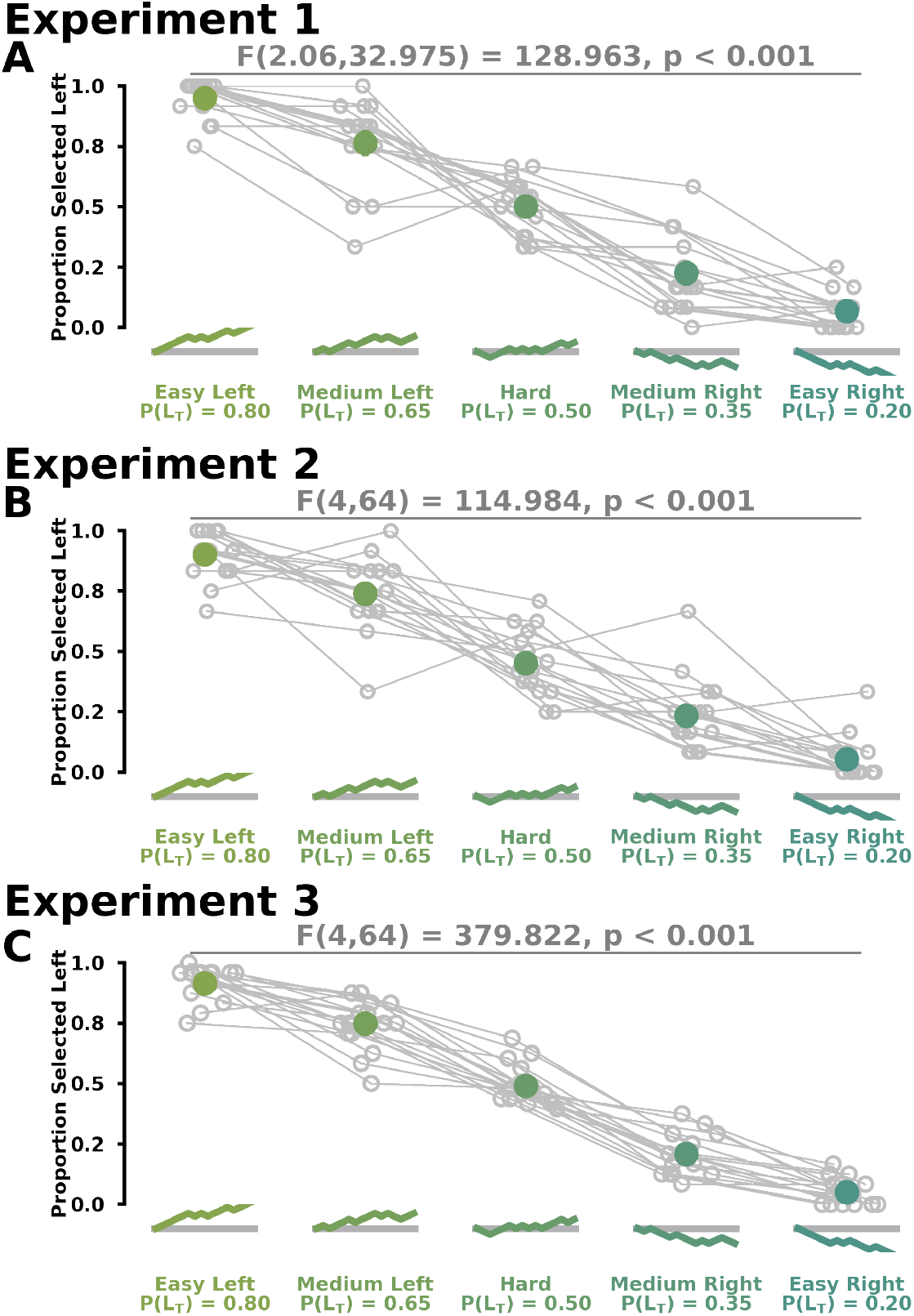
Group Selection Rate Behaviour for Pseudorandom Token Patterns. Proportion of Left Selections (y-axis) across pseudorandom token patterns (x-axis) in **A) Experiment 1, B) Experiment 2**, and **C) Experiment 3**. Inset text shows significant effects from an rmANOVA.

## Supplementary D

### Selection Rate Behaviour

#### Bias Pattern Selection Rates

**Figure SD1:**
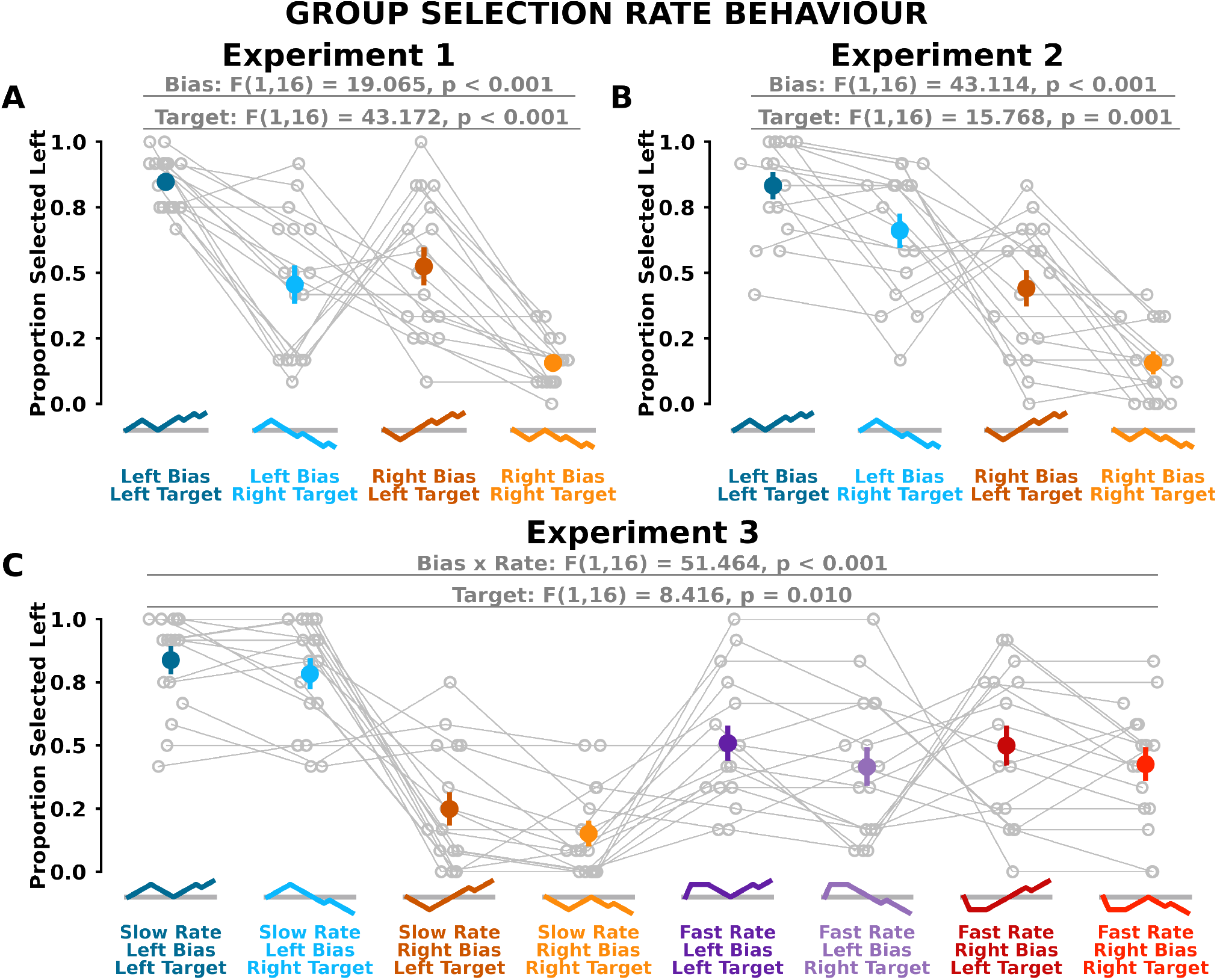
Selection Rates for Bias Token Patterns. Proportion of left target selection trials (y-axis) across bias token patterns (x-axis) in **A) Experiment 1, B) Experiment 2**, and **C) Experiment 3**. Inset text shows significant effects from ANOVA analysis.

#### Bias Pattern Selection Rate Over Time

**Figure SD2:**
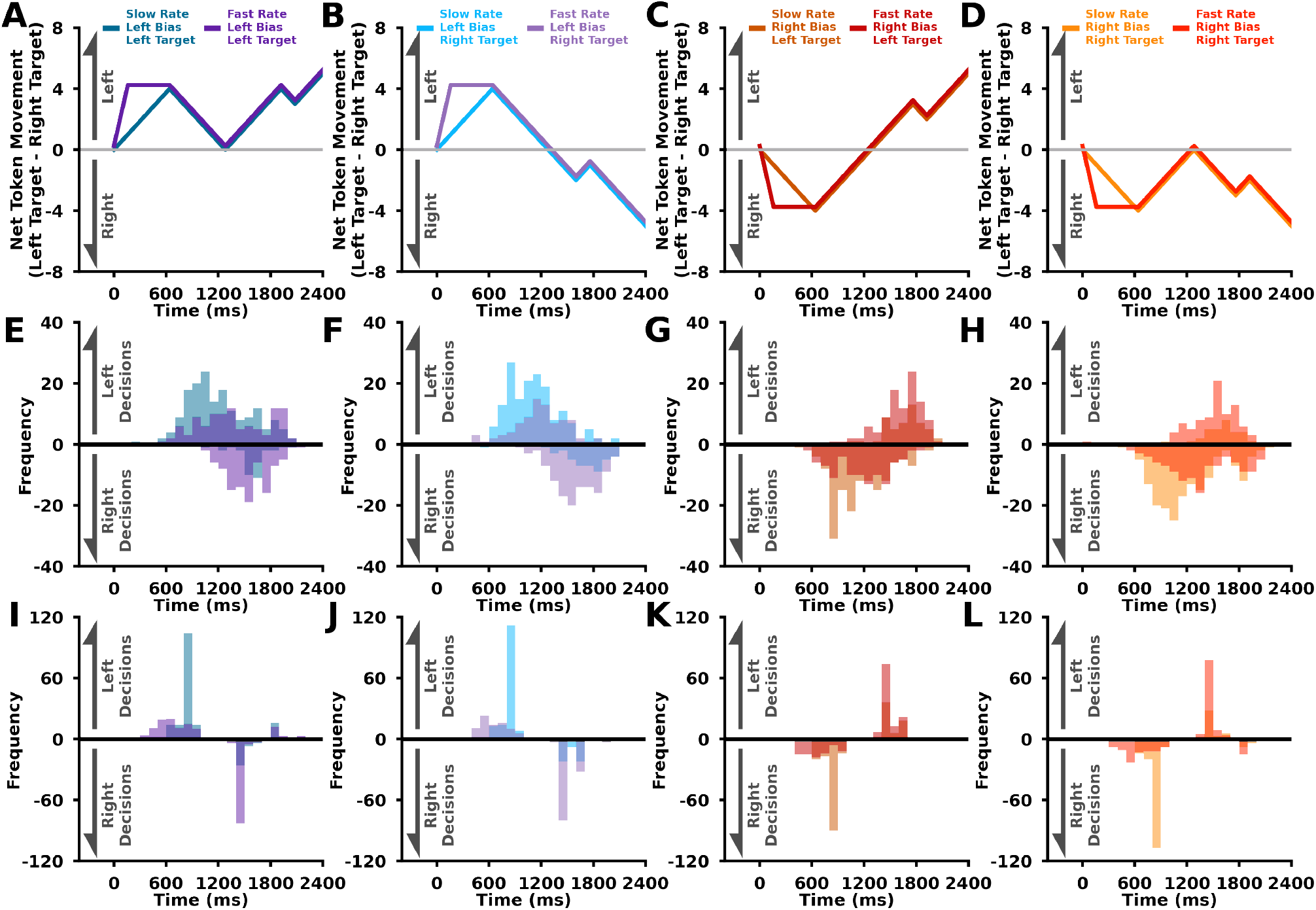
Selection Rate Distributions for Rate Bias Token Patterns in Experiment 3. **A-D)** Net token movement (left target - right target; y-axis) over time (x-axis). Each plot shows the slow rate and fast rate token patterns for the same bias and final target. **E-H)** Histogram of group behaviour estimated decision time (x-axis) for left and right decisions. **I-L)** Trueblood (2001) model with novel evidence using best-fit parameters. Histogram of model predicted estimated decision time (x-axis) for left and right decisions. Histogram colors are representative of the token patterns in the plot directly above. Positive and negative histograms represent left decisions and right decisions respectively.

## Supplementary E

### Modelling Methods

#### Decision-Making and Movement Model

Here we compared five types of decision-making models which predicted target selections and decision times given sensory information or evidence for a given target. We also compared two types of evidence, current or novel, as input into the decision-making models.

### Evidence

As inputs to the decision-making models, we used novel evidence or current evidence. Evidence is based on the correct probability (*p*(*L* | *N*_*L*_, *N*_*C*_, *N*_*R*_)) for the left target

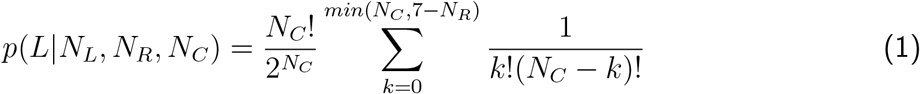

given the number of tokens in the left target (*N*_*L*_), number of tokens between the targets (*N*_*R*_), and the number of tokens in the right target (*N*_*R*_; Equation 1), and ! represents a factorial.^14^ Current evidence (*E*_*curr*_(*t*)) is defined as the correct probability at the current time with added sensory noise

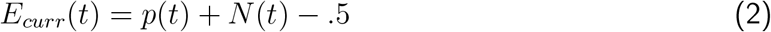

where *t* is time, *N* (*t*) is sensory noise modelled with a gaussian that has a zero mean and a standard deviation (*σ*_*ev*_). Novel evidence (*E*_*novel*_(*t*)) is defined as the rate of change (*d/dt*) of the correct probability with added sensory noise (Equation 3).

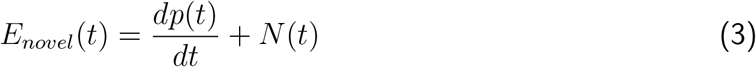

### Decision Making Models

In our decision making models, we simulate a decision variable that interprets the evidence used to make a decision. We define a decision as the time when the decision variable crosses a threshold of +1.0 for a left target decision or -1.0 for a right target decision.

#### Drift-diffusion model.^17,18^

The rate of change of the decision variable (DV) is equal to a gain (g) multiplied by the evidence.

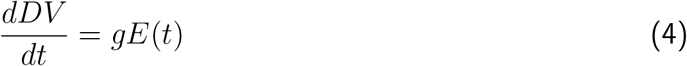

#### Drift diffusion model with leak^19,20,14^

The rate of change of the decision variable is similar to equation 4, but with a leak (L) term that represents forgetting.

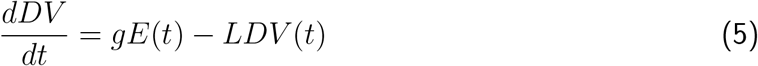

#### Trueblood model (2021).^24^

The rate of change of the decision variable is a function of urgency (k) and leak (L; eq. 6). It can be seen that the urgency term k is scaled by time (t) to influence the weightings of incoming evidence (second term on right side of the equation) and previously accumulated evidence (first term on right side of the equation).

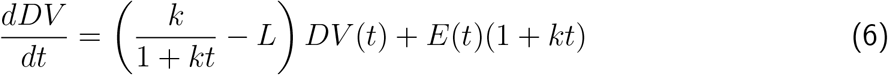

#### Urgency-gating model.^14^

The decision variable is equal to the evidence scaled by a temporally increasing urgency signal (*U* (*t*)); eq. 7). The urgency signal is a scalar (g) multiplied by the current time (eq. 8).

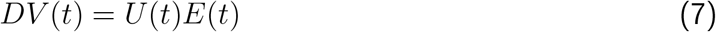

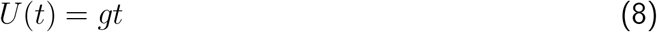

#### Urgency-gating model with a low-pass filter.^14,15^

The decision variable is equal to an estimate of the evidence (*E*_*est*_) scaled by a temporally increasing urgency signal (*U* (*t*); eq. 9). The estimate of evidence is a low-pass filtered version of the incoming evidence (eq.10). The urgency signal is a scalar (g) multiplied by the current time (eq. 11).

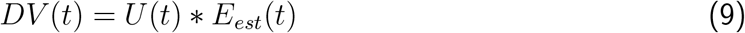

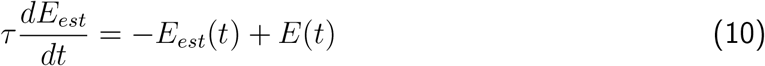

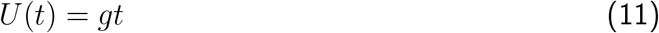

We simulated each trial until either the decision variable crossed a decision threshold or the trial deadline (2400 ms). Evidence was input into the decision making models with a 200ms delay. We used a time step of 1 ms for all decision-making simulations.

### Movement Model

Here we use a linear quadratic gaussian optimal feedback controller,^25,26,27,28,29,30^ which used the decision variable from the Trueblood model to weight potential goals. The dynamics of the hand are

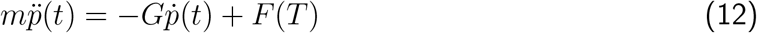

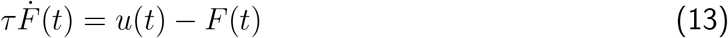

where m is mass (1 kg), p(t) is the position of the hand, and G is the viscous constant 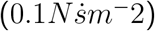. u(t) is a control signal (e.g., muscle activity). F(t) represents internal forces (e.g., muscle force) that move the hand. *τ* is a low-pass filter time constant (40 ms) that approximates the rate of internal forces given some control signal.^70,28^ Single and double dots refer to single and double differentiation.

We transformed the decision variable into a weighting (*α*, (eq. 12)) for each option using a logistic function

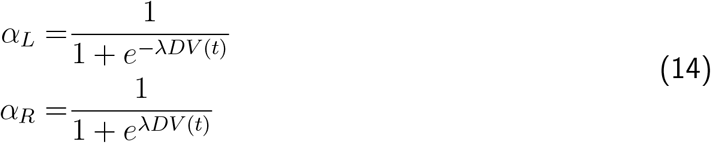

where *λ* is the steepness parameter and t is the current time step.

We also define the location of the reaching target as (*p*_*G*_) where alpha is the weighting for each target from eq. 12. *p*_*x,L*_ and *p*_*y,L*_ correspond to the forward and lateral position of the left target. *p*_*x,R*_ and *p*_*y,R*_ correspond to the forward and lateral position of the right target.

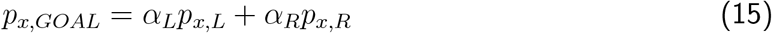

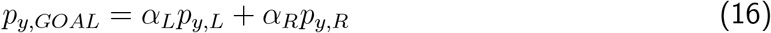

We combine the states into a state vector (x) in eq. 17.

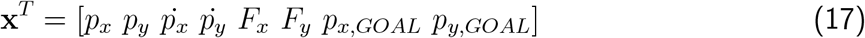

The dynamics of the system is then discretized with added state noise in eq. 18. The covariance matrix of the state noise is a matrix with [0,0,0,0,1e-3,1e-3] on the diagonals and zeros elsewhere.

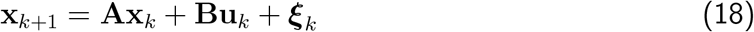

We define a standard quadratic cost function with a (**Q**_**N**_) terminal cost, running state cost (**Q**), and control costs (**R**). Note that **Q** was constant for all time steps. N is the total number of time steps in the trial (240).

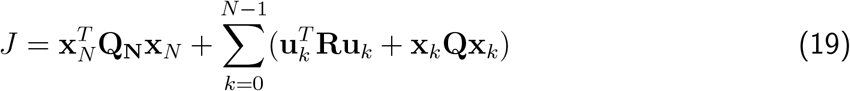

We define **Q** and **Q**_**N**_ such that

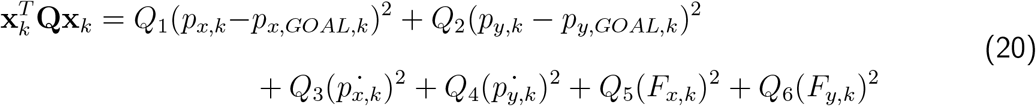

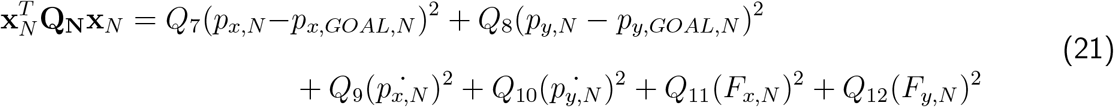

The sensory feedback signal (**x**) is equal to the current state with added sensory noise (*η*; eq. 20).

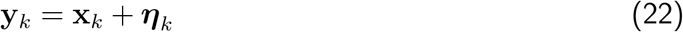

The covariance matrix for the sensory noise is a matrix with [1e-3,1e-3,1e-3,1e-3,1e-3,1e-3] on the diagonals and zeros elsewhere.

We used a Kalman filter (K) to estimate the current state (eq. 23).^27,28^ We used the standard calculation of the Kalman filter.^25,26,27,28,29,30^ Note we did not consider signal dependent noise, which would not have a large influence on our results given the very large target sizes.

Here, 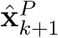 is the prior belief of the next state, based on 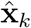 and the control signal *u*_*k*_. The state estimate 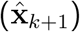 is dependent on the prior 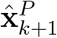, the kalman filter (**K**), and the sensory feedback (**y**_*k*+1_).

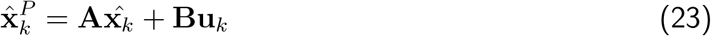

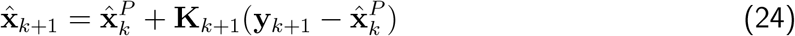

We solved for an optimal feedback policy (**L**) as a function of the cost function and given dynamics^25,26,27,28,29,30^ using the Ricatti equations. We use the optimal feedback policy to calculate the current optimal control signal. On each timestep, the optimal feedback controller generates an optimal control signal (**u**_*k*_) that feeds into the dynamics (eq. 18) as

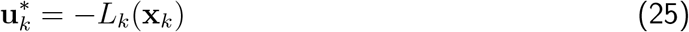

Each model simulation runs until the point mass enters one of the two targets (see Experiment 2 Methods) or runs out of time (2400 ms). We simulated movements with a time step of 10 ms.

### Decision-Making Model Fitting Procedure

In total we fit and tested ten models (five decision-making models x two types of evidence). We used the same fitting procedure for each model. Model fitting was performed using the powell algorithm in the Minimize function from the Scipy Python library. We fit each experiment separately. For each experiment, we only fit the model to the behaviour during the bias token patterns.

For each model, we simulated 500 trials for each bias token pattern. We then calculated the mean decision time and selection rate for each bias token pattern. The loss function was defined using the decision time and selection rate. For the decision time, we calculated the difference between model mean decision times and data mean decision times, then normalized by 2400 ms. We then the absolute value of this normalized error. For the selection rate, we calculated the difference between model mean selection rate and data mean selection rate, then normalized by 100%. We then took the absolute value of this normalized error. To calculate the final loss, we summed across token patterns (Y) for both decision times (DT) and selection rates (SR). When fitting Experiment 3, we also considered the average difference of decision times between the slow and fast rate token patterns.

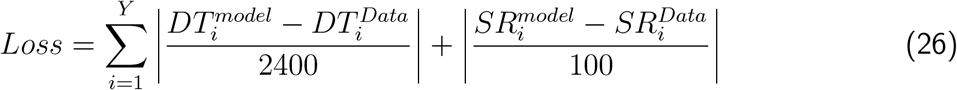

For the fitting procedure we first began with a warm-start procedure.^67,71^ First, we fit the model 1,000 times using random initial parameter guesses. From these fits we then selected the model parameters that resulted in the lowest loss. The lowest loss parameters were then used as an initial guess for a bootstrapping procedure (10,000 iterations) to find the 95% confidence interval of each parameter given the data. In each bootstrap iteration, we resampled with replacement the decision time and selection rate from the data. The mean decision times and selection rates of the resampled data were then used to determine model loss for each bootstrap.

### Movement Model Fitting Procedure

For the movement model, we fit the terminal state costs parameters (**Q**_**n**_), running state cost parameters (**Q**), running energetic costs (**R**), and steepness parameter (*λ*) of the logistic function. We fit Experiment 2 and 3 simultaneously.

We first simulated decision variables using the Trueblood Model with novel evidence. We used the median model parameters from the boot-strapping procedure for each experiment. Model fitting was performed using the powell algorithm in the Minimize function from the Scipy Python library. We simulated 500 trials for each bias token pattern. We calculated the mean trajectory for each token pattern from the simulated trials. The loss function was defined as the squared error between the group mean trajectory and the simulated mean trajectory. We used the model parameters that resulted in the lowest loss.

### Haith and Wong

To simulate the Wong and Haith (2017) study, we used our decision-making and movement model.^3^ We selected parameters that qualitatively resulted in proportions of intermediate movements and trajectories that matched the experimental behaviour. Importantly, we defined the urgency parameter (k; see eq. 6) as a function of the current task condition (eq. 23). As suggested in Carland 2019, we used an urgency signal (k) that was a function of reward, energetic cost, and time.^31^

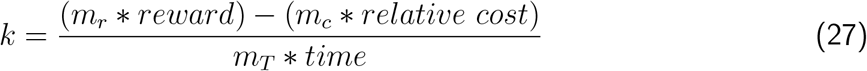

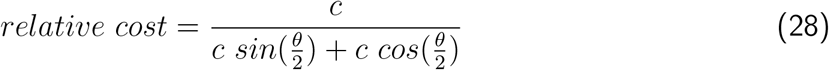

where *m*_*r*_ is the weighting on reward (8), reward was the value of success (1), *m*_*c*_ was the weighting of a direct reach relative to a intermediate reach (6), *m*_*T*_ is the weighting on relative time (0.002), and time is the time participants had to reach a target. Here we considered the relative energetic costs of reaching a shorter distance directly from the start position to a target (e.g., the hypotenuse of a right angle triangle), compared to travelling an overall further distance by first reaching between the targets (e.g., adjacent side of a triangle) and then to one of the targets (e.g., along the adjacent and then opposite side of a triangle). Specifically, we calculated the ratio between the hypotenuse (c = 20) and the distance of travelling along the adjacent and opposite sides of the corresponding right triangle, given the angular distance of the targets about the start position (*θ*: 15, 30, 45, 60). Similarly, as a proxy for time, we approximated the time participants had to reach a target (slow: 1000ms, fast: 500ms) given the experimentally imposed slow and fast hand movement criteria.

### Goal Averaged Single Flexible Plan versus Averaged Parallel Motor Plans

Here we define the discrete state dynamics (eq. 29) and the cost function (eq. 30) as the same as above in the modelling methods.

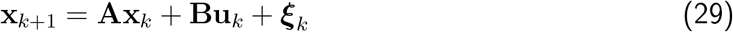

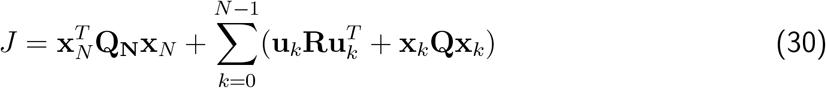

We solve for the optimal feedback control policy (**L**_*k*_) using the riccati equation. The optimal control signal 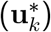 is defined as

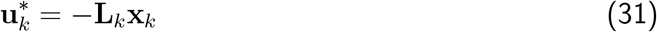

where **x**_*k*_ is the current state. Given the assumptions of our cost function in eq. 19-21, we can consider the optimal feedback control signal as equal to

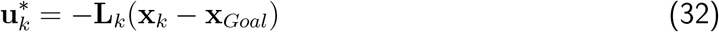

To match our experimental design, we consider the goal to be a weighted average of the two potential targets given the current decision variable (see eq. 15, 16). We rewrite eq. 32 as

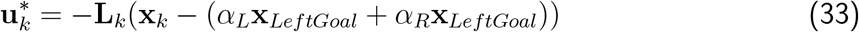

This can be thought as a single flexible control policy to a goal averaged target.

The weighting terms are calculated from a logistic function (eq. 14) with bounds 0 and 1, and thus *α*_*L*_ + *α*_*R*_ = 1. We can then expand eq. 33 as

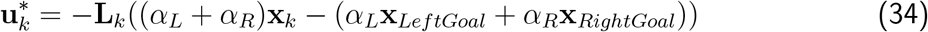

We further rearrange eq 34 as

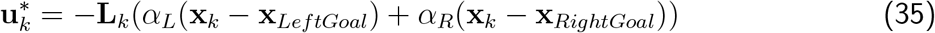

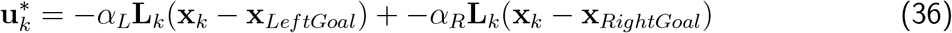

It can now be seen that 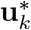 is the weighted sum of two optimal feedback control policies for each potential goal. In other words, this can be considered as the average of parallel flexible control policies. It is important to note that this holds given the assumption that the dynamics and costs are the same between the possible targets. Given the assumptions above, parallel averaged flexible control policies and a single flexible control policy to an averaged goal are not dissociable.

### Modelling Outcomes

**Figure SE1:**
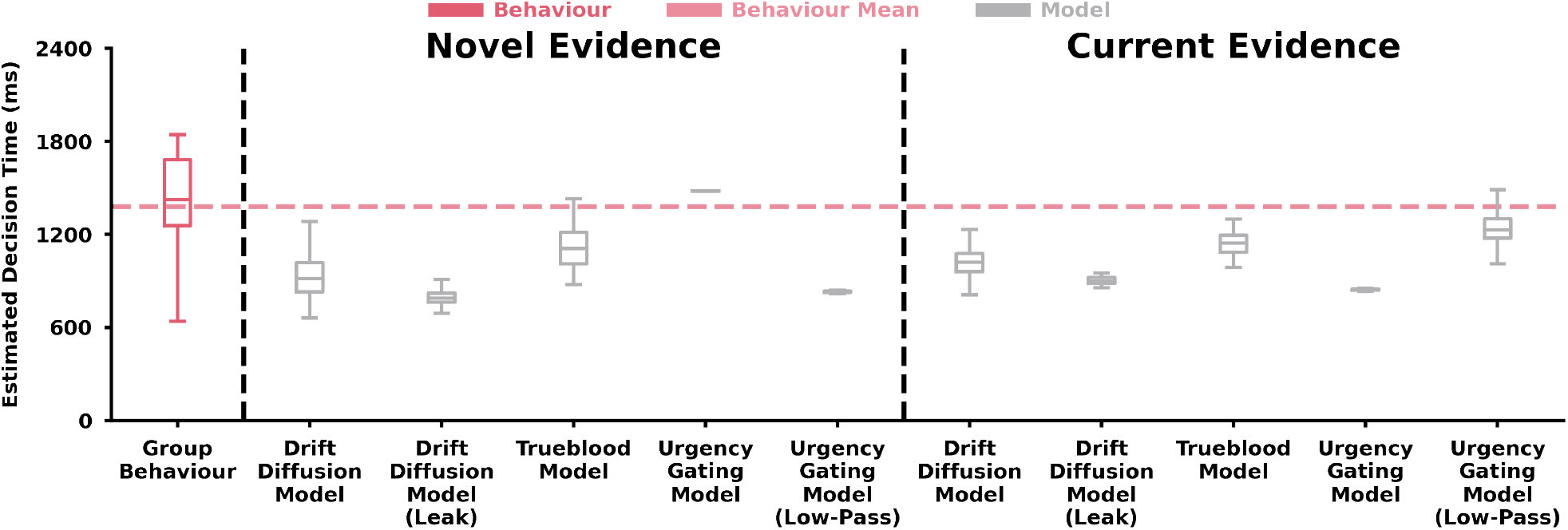
Experiment 1 Best-Fit Parameter Model Simulation. **Experiment 2** Estimated Decision Time (y-axis) for Group Behaviour and Decision-Making Models (x-axis). Group participant estimated decision times are shown for bias token patterns (dark pink). Best-fit model simulations of decision times are shown for bias token patterns (light grey). Dashed light pink line represents mean estimated decision time behaviour. Box and whisker plots show 25%, 50% and 75% quartiles. Inset labels represent models simulated with novel sensory evidence or current sensory evidence.

**Figure SE2:**
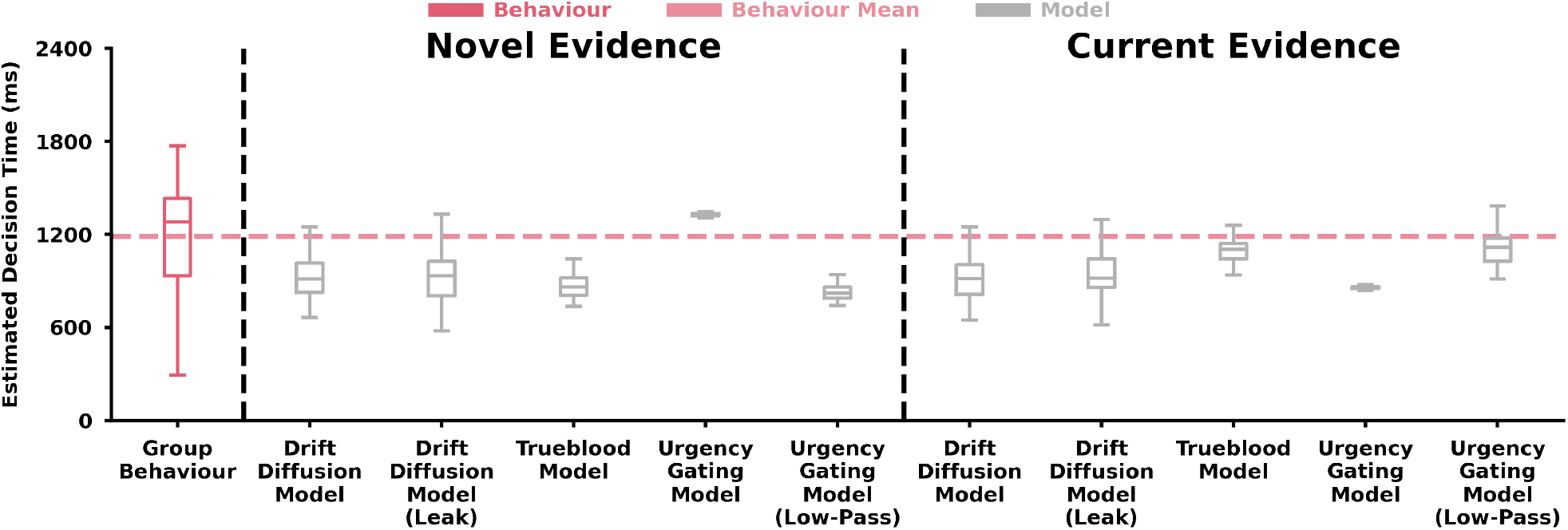
Experiment 2 Best-Fit Parameter Model Simulation. **Experiment 2** Estimated Decision Time (y-axis) for Group Behaviour and Decision-Making Models (x-axis). Group participant estimated decision times are shown for bias token patterns (dark pink). Best-fit model simulations of decision times are shown for bias token patterns (light grey). Dashed light pink line represents mean estimated decision time behaviour. Box and whisker plots show 25%, 50% and 75% quartiles. Inset labels represent models simulated with novel sensory evidence or current sensory evidence.

**Figure SE3:**
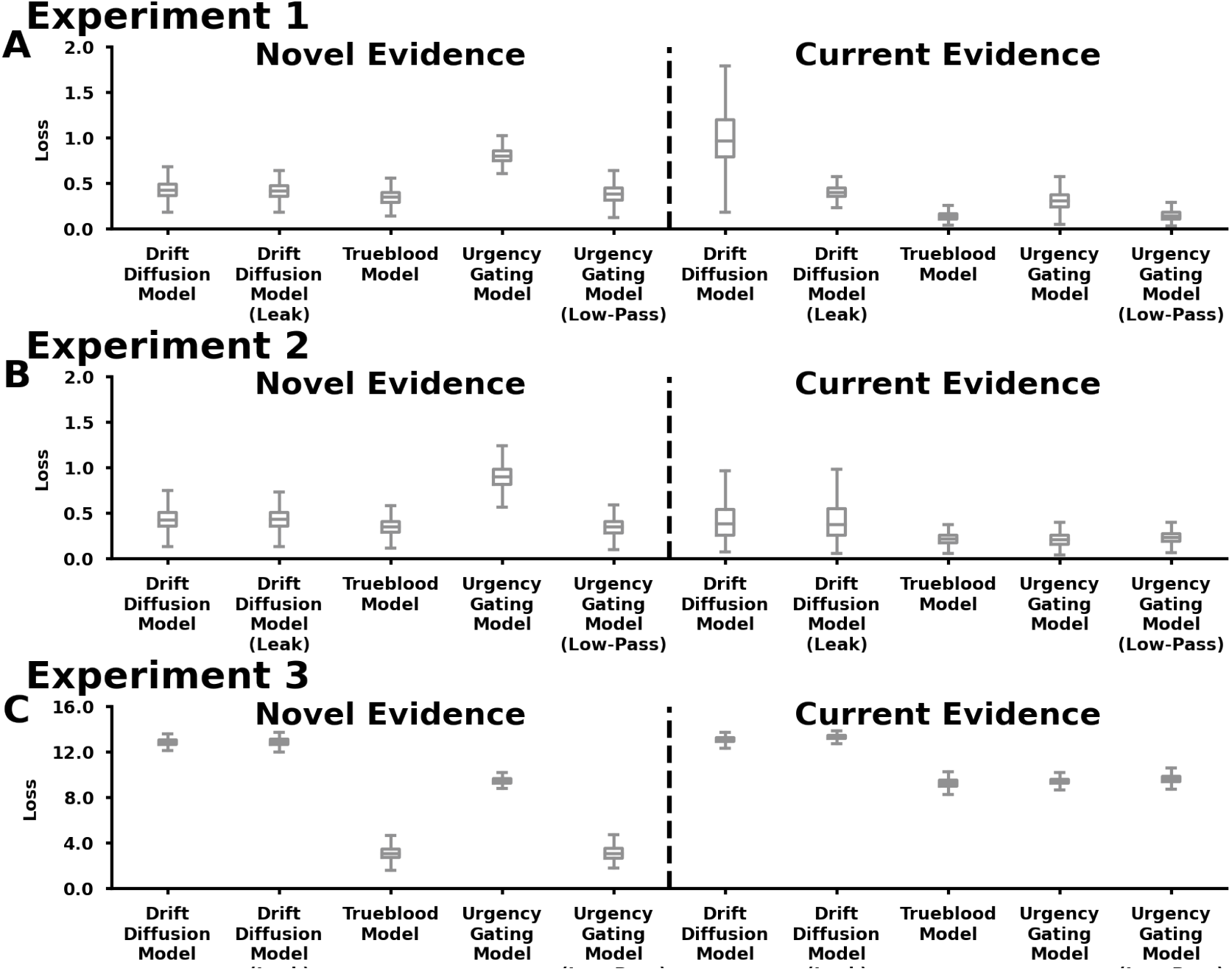
Model Loss. Bootstrapped loss values (y-axis) across decision making models (x-axis) in **A) Experiment 1, B) Experiment 2**, and **C) Experiment 3**. Inset labels represent models simulated with novel sensory evidence or current sensory evidence.

**Table SE1:** Experiment 1: Model Parameters.

|  | Sensory Evidence | Parameters |  |  |
| --- | --- | --- | --- | --- |
| Drift Diffusion Model | Novel | Gain | | Noise ($\sigma$) |
|  |  | 2.91 [2.82, 3.11] |  | 2.28 [1.47, 4.95] |
| Drift Diffusion Model With Leak | Novel | Gain | Leak | Noise ($\sigma$) |
|  |  | 4.08 [4.01, 4.16] | 1.58 [1.48, 1.72] | 0.76 [0.31, 1.97] |
| Trueblood Model | Novel | Urgency | Leak | Noise ($\sigma$) |
|  |  | 1.79 [1.34, 2.53] | 0.08 [0.00, 0.35] | 4.83 [3.04, 6.29] |
| Urgency-Gating Model | Novel | Urgency | | Noise ($\sigma$) |
|  |  | 0.01 [0.01, 0.01] |  | 8.51 [3.44, 15.56] |
| Urgency-Gating Model with Low-Pass Filter | Novel | Urgency | Tau | Noise ($\sigma$) |
|  |  | 17.61 [17.04, 17.84] | 4.24 [4.17, 4.35] | 0.42 [0.12, 1.70] |
| Drift Diffusion Model | Current | Gain | | Noise ($\sigma$) |
|  |  | 6.19 [5.56, 6.58] |  | 2.24 [0.23, 4.10] |
| Drift Diffusion Model With Leak | Current | Gain | Leak | Noise ($\sigma$) |
|  |  | 16.76 [15.96, 17.72] | 4.56 [4.37, 4.89] | 0.94 [0.81, 1.14] |
| Trueblood Model | Current | Urgency | Leak | Noise ($\sigma$) |
|  |  | 16.10 [14.33, 18.33] | 9.82 [8.82, 11.75] | 2.60 [2.46, 2.89] |
| Urgency-Gating Model | Current | Urgency | | Noise ($\sigma$) |
|  |  | 2.50 [2.20, 2.58] |  | 0.06 [0.06, 0.08] |
| Urgency-Gating Model with Low-Pass Filter | Current | Urgency | Tau | Noise ($\sigma$) |
|  |  | 1.42 [1.26, 1.72] | 0.12 [0.11, 0.14] | 3.69 [3.41, 4.05] |

**Table SE2:** Experiment 2: Model Parameters.

**Table SE1: Experiment 1: Model Parameters.**
|  | Sensory Evidence | Parameters |  |  |
| --- | --- | --- | --- | --- |
| Drift Diffusion Model | Novel | Gain | | Noise ($\sigma$) |
|  |  | 2.83 [2.67, 3.22] |  | 5.93 [2.32, 7.78] |
| Drift Diffusion Model With Leak | Novel | Gain | Leak | Noise ($\sigma$) |
|  |  | 3.13 [2.94, 3.60] | 0.46 [0.12, 1.12] | 5.44 [2.43, 7.19] |
| Trueblood Model | Novel | Urgency | Leak | Noise ($\sigma$) |
|  |  | 2.64 [2.11, 3.13] | 0.14 [0.01, 0.46] | 3.49 [1.60, 5.13] |
| Urgency-Gating Model | Novel | Urgency | | Noise ($\sigma$) |
|  |  | 0.01 [0.01, 0.01] |  | 10.19 [5.01, 16.79] |
| Urgency-Gating Model with Low-Pass Filter | Novel | Urgency | Tau | Noise ($\sigma$) |
|  |  | 22.45 [19.58, 25.09] | 5.65 [5.21, 6.45] | 2.75 [0.63, 4.53] |
| Drift Diffusion Model | Current | Gain | | Noise ($\sigma$) |
|  |  | 5.01 [4.70, 5.86] |  | 5.07 [3.66, 5.76] |
| Drift Diffusion Model With Leak | Current | Gain | Leak | Noise ($\sigma$) |
|  |  | 5.75 [5.38, 6.87] | 0.60 [0.33, 1.58] | 4.89 [3.85, 5.53] |
| Trueblood Model | Current | Urgency | Leak | Noise ($\sigma$) |
|  |  | 13.14 [11.24, 15.78] | 9.35 [7.32, 11.88] | 3.92 [3.46, 4.63] |
| Urgency-Gating Model | Current | Urgency | | Noise ($\sigma$) |
|  |  | 1.98 [1.88, 2.06] |  | 0.11 [0.11, 0.11] |
| Urgency-Gating Model with Low-Pass Filter | Current | Urgency | Tau | Noise ($\sigma$) |
|  |  | 9.56 [7.17, 13.64] | 2.16 [1.81, 2.67] | 6.39 [4.98, 7.97] |

**Table SE3:** Experiment 3: Model Parameters.

**Table SE2: Experiment 2: Model Parameters.**
|  | Sensory Evidence | Parameters (median [95% CI]) |  |  |
| --- | --- | --- | --- | --- |
| Drift Diffusion Model | Novel | Gain<br>19.47 [15.38, 25.15] | | Noise ($\sigma$)<br>2.74 [1.70, 5.09] |
| Drift Diffusion Model With Leak | Novel | Gain<br>19.21 [15.25, 24.48] | Leak<br>5.83 [1.16, 11.98] | Noise ($\sigma$)<br>3.62 [2.40, 6.01] |
| Trueblood Model | Novel | Urgency<br>2.80 [2.24, 3.09] | Leak<br>0.92 [0.70, 1.16] | Noise ($\sigma$)<br>2.63 [2.21, 3.52] |
| Urgency-Gating Model | Novel | Urgency<br>0.01 [0.00, 0.01] | | Noise ($\sigma$)<br>5.22 [0.82, 12.71] |
| Urgency-Gating Model with Low-Pass Filter | Novel | Urgency<br>4.94 [4.21, 5.68] | Tau<br>1.39 [1.13, 1.74] | Noise ($\sigma$)<br>2.94 [2.13, 4.28] |
| Drift Diffusion Model | Current | Gain<br>2.18 [1.59, 4.05] | | Noise ($\sigma$)<br>21.35 [16.49, 26.45] |
| Drift Diffusion Model With Leak | Current | Gain<br>15.28 [13.83, 17.66] | Leak<br>186.87 [181.14, 194.57] | Noise ($\sigma$)<br>13.24 [11.32, 15.62] |
| Trueblood Model | Current | Urgency<br>2.15 [1.57, 3.03] | Leak<br>12.97 [8.78, 21.28] | Noise ($\sigma$)<br>20.73 [16.17, 25.44] |
| Urgency-Gating Model | Current | Urgency<br>0.92 [0.80, 1.00] | | Noise ($\sigma$)<br>0.23 [0.20, 0.28] |
| Urgency-Gating Model with Low-Pass Filter | Current | Urgency<br>0.15 [0.11, 0.20] | Tau<br>0.01 [0.01, 0.03] | Noise ($\sigma$)<br>11.54 [8.03, 15.80] |

**Table SE4:** Movement Model Model Parameters.

**Table SE3: Experiment 3: Model Parameters.**
| Name | Value |
| --- | --- |
| $Q_1$ | 3.08e-05 |
| $Q_2$ | 3.32e-01 |
| $Q_3$ | 7.40e-05 |
| $Q_4$ | 1.62 |
| $Q_5$ | 8.57e-03 |
| $Q_6$ | 8.76e-03 |
| $Q_7$ | 5.55e+02 |
| $Q_8$ | 1.48e+03 |
| $Q_9$ | 4.09 |
| $Q_{10}$ | 1.56e+02 |
| $Q_{11}$ | 1.47e+02 |
| $Q_{12}$ | 3.21e+01 |
| $R$ | 1.87e-07 |
| $\lambda$ | 2.58 |

